# Human iPSC-based neurodevelopmental models of globoid cell leukodystrophy uncover patient- and cell type-specific disease phenotypes

**DOI:** 10.1101/2020.03.13.990176

**Authors:** Elisabeth Mangiameli, Anna Cecchele, Francesco Morena, Francesca Sanvito, Vittoria Matafora, Angela Cattaneo, Lucrezia Della Volpe, Daniela Gnani, Marianna Paulis, Lucia Susani, Sabata Martino, Raffaella Di Micco, Angela Bachi, Angela Gritti

**Affiliations:** San Raffaele Telethon Institute for Gene Therapy (SR-Tiget), IRCCS San Raffaele Scientific Institute, Via Olgettina 60, 20132 Milan, Italy; Department of Chemistry, Biology and Biotechnology, University of Perugia, Via del Giochetto, 06123 Perugia, Italy; Pathology Unit, IRCCS San Raffaele Scientific Institute, Via Olgettina 60, 20132 Milan, Italy; IFOM-FIRC Institute of Molecular Oncology, Via Adamello 16, 20139 Milan, Italy; Humanitas Clinical and Research Center-IRCCS, Rozzano, Milan, Italy; National Research Council (CNR)-IRGB/UOS of Milan, Milan, Italy

**Keywords:** Pluripotent stem cells, leukodystrophy, Krabbe disease, gene therapy, neural progenitors, oligodendrocytes, senescence

## Abstract

Globoid Cell Leukodystrophy (GLD, or Krabbe disease) is a rare lysosomal storage disease caused by inherited deficiency of β-galactocerebrosidase (GALC). The build-up of galactosylsphingosine (psychosine) and other undegraded galactosylsphingolipids in the nervous system causes severe demyelination and neurodegeneration. The molecular mechanisms of GLD are poorly elucidated in neural cells and whether murine systems recapitulate critical aspects of the human disease is still to be defined.

Here, we established a collection of GLD patient-specific induced pluripotent stem cell (iPSC) lines. We differentiated iPSCs from two patients (bearing different disease-causing mutations) into neural progenitors cells (NPCs) and their neuronal/glial progeny, assessing the impact of GALC deficiency and lentiviral vector-mediated GALC rescue/overexpression by means of phenotypic, biochemical, molecular, and lipidomic analysis. We show a progressive increase of psychosine during the differentiation of GLD NPCs to neurons and glia. We report an early and persistent impairment of oligodendroglial and neuronal differentiation in GLD cultures, with peculiar differences observed in the two GLD lines. GLD cells display a global unbalance of lipid composition during the iPSC to neural differentiation and early activation of cellular senescence, depending on the disease-causing mutation. Restoration of GALC activity normalizes the primary pathological hallmarks and partially rescues the differentiation program of GLD NPCs.

Our results suggest that multiple mechanisms besides psychosine toxicity concur to CNS pathology in GLD and highlight the need of a timely regulated GALC expression for proper lineage commitment and differentiation of human NPCs. These findings have important implications for establishing tailored gene therapy strategies to enhance disease correction in GLD.

## Introduction

Globoid Cell Leukodystrophy (GLD, or Krabbe disease) is a rare lysosomal storage disorder (LSD) caused by inherited deficiency of β-galactocerebrosidase (GALC), a key enzyme in the catabolism of galactosphingolipids, e.g. galactosylceramide and galactosylsphingosine (psychosine) (Suzuki and Suzuki, 1970). The infantile forms are characterized by relentless demyelination and neurodegeneration of the central and peripheral nervous system (CNS, PNS). Allogeneic hematopoietic stem/progenitor cell (HSPC) transplant - the only therapeutic option for GLD infants - provides a functional GALC enzyme to the affected organs through the engraftment of HSPC-derived myeloid progeny. However, it fails to fully correct the severe CNS pathology, even if performed at the asymptomatic stage (Allewelt et al., 2016), likely because of the insufficient GALC supply provided by HSPC-derived microglia in the brain and/or modest metabolic correction of endogenous cells. *In vivo* and *ex vivo* gene therapy (GT) approaches using lentiviral vectors (LV) and adeno-associated vectors (AAV) benefit GLD animal models (Bradbury et al., 2018; Gentner et al., 2010; Lattanzi et al., 2010, 2014; Lin et al., 2005; Meneghini et al., 2016; Rafi et al., 2015; Ricca et al., 2015; Ungari et al., 2015). The advantage of *ex vivo* GT with LV-modified HSPCs in infants affected by metachromatic leukodystrophy (MLD) – a similar LSD (Biffi et al., 2013; Sessa et al., 2016), suggests that a similar approach might be translated to GLD. Still, the obstacles in reaching safe, widespread and stable expression of therapeutic GALC levels in the human CNS limits the clinical development of these GT strategies to GLD.

Studies in GLD mice suggest that a modest accumulation of psychosine impairs CNS and PNS development from neural progenitor cells (NPCs) (Castelvetri et al., 2011; Santambrogio et al., 2012; Teixeira et al., 2014). Then, psychosine accumulates in GLD neurons and glial cells until reaching a threshold that triggers cytotoxicity. Besides the effects exerted on myelin and axons, psychosine and other substrates in the GALC-related sphingolipid pathway (Hannun and Obeid, 2017) directly or indirectly affect glial and neuronal cell homeostasis interfering with lipid content and distribution (Giri et al., 2006; White et al., 2009), altering proteostasis network (Pellegrini et al., 2019), promoting apoptosis (Jatana et al., 2002), lysosomal dysfunction (Folts et al., 2016; Samie and Xu, 2014), and autophagy (Del Grosso et al., 2019). Growing evidence points to a previous underestimated association between lysosomal dysfunction, protein misfolding/aggregation, lipid unbalance, and senescence in multiple organisms and cell types (Buratta et al., 2017; Irahara-Miyana et al., 2018; Lee et al., 2014; Lizardo et al., 2017; Smith et al., 2014; Trayssac et al., 2018). Also, recent observations implicate NPCs and oligodendroglial progenitor cells (OPCs) senescence in the pathogenesis of neurodegenerative disorders with protein aggregation (Zhang et al., 2019) and neuroinflammatory demyelinating disorders (Nicaise et al., 2019), providing a strong rationale for investigating the potential implication of these processes in GLD. Altogether, these complex and yet poorly elucidated events may contribute to the multifaceted aspects of GLD pathology and explain the modest efficacy of conventional or experimental therapies in correcting CNS and PNS damage in GLD animal models and patients (Allewelt et al., 2016; Hawkins-Salsbury et al., 2015; Rafi et al., 2015; Ricca et al., 2015).

More than 100 disease-causing mutations in the human *GALC* gene have been described (Wenger and Luzi, 2014), among which a 30-kb deletion accounts for 40% of pathogenic alleles in the Caucasian population (Rafi et al., 1995). The occurrence of this large gene deletion in homozygosis is associated to the early-onset and most severe disease forms. The genotype-phenotype correlation is less clear for the many missense/nonsense mutations occurring in homozygosis or compound heterozygosis, which may be associated to variable residual GALC activity and severe to mild disease variants (Duffner et al., 2011, 2012; De Gasperi et al., 1996, 1999; Jalal et al., 2012; Tappino et al., 2010). Misfolded GALC proteins evoke different Unfolded Protein Response (UPR) activation (Irahara-Miyana et al., 2018) and show mutation-specific alteration of trafficking to lysosomes and processing (Shin et al., 2016), suggesting the occurrence of pathological cascades that are independent from or act in synergy with primary lipid storage in determining the complex GLD phenotype. Thus, studying the mechanisms linking the primary mutation-specific event(s) to progressive cell dysfunction and death is relevant and requires appropriate experimental models.

Murine, canine and non-human primate models of GLD (Wenger, 2000) have been used to test the safety and efficacy of novel therapies; nevertheless, they fail to fully recapitulate the spectrum of pathological manifestations observed in patients. Also, the most used mouse model (Suzuki, 1983) and non-human primate model of GLD (Borda et al., 2008; Luzi et al., 1997) carry spontaneous mutations that are not found in humans. Conventional GLD human cellular models include patient-specific fibroblasts (Berardi et al., 2014; Gama Sosa et al., 1996; Rafi et al., 1996; Ribbens et al., 2013; Spratley et al., 2016), human hematopoietic cells (Martino et al., 2009), or epithelial cell lines with induced GALC mutations (Lee et al., 2010; Shin et al., 2016) that hardly recapitulate the metabolic and functional features of NPCs and their differentiated progeny. Thus, there is a strong need for human-derived experimental models to explore GLD pathogenesis and test correction strategies.

Human induced pluripotent stem cells (iPSCs) (Yamanaka, 2007) offer a unique tool to analyze the molecular consequences of pathogenic mutations in the context of the genetic background of individual patients. The differentiation of iPSCs in neural cells has boosted CNS disease modelling and therapeutic screening (Shi et al., 2017). Human iPSCs are available from a variety of LSDs (Huang et al., 2012) but there are no reports describing GLD patient-derived iPSCs.

Here, we established GLD patient-specific iPSC lines as a reliable human model for elucidating GLD pathogenesis and testing the biological efficacy of gene therapy in relevant neural cell types. To this end, we differentiated iPSCs into NPCs and differentiated cells (oligodendrocytes, neurons, and astrocytes) and monitored the appearance/progression of cell type- and patient-specific primary and secondary defects in GLD cells. Finally, we assessed the impact of GALC reconstitution/overexpression (achieved by lentiviral-mediated gene transfer) in reverting the pathological phenotype and its potential impact on the biology of human NPCs and progeny.

## Results

### GLD iPSCs show undetectable GALC activity and psychosine storage but display normal pluripotency features

We generated iPSCs from fibroblasts of 5 GLD patients with distinct biallelic mutations in the *GALC* gene (homozygous or compound heterozygous) resulting in the infantile form of the disease. Control iPSC lines were generated from fibroblasts of two unrelated normal donors (ND1, adult; ND2, neonatal) and one non-affected relative (ND3, parent of GLD1)(**Table S1**). GALC activity in ND fibroblasts was 2.42 nmol*h/mg (ND1), 3.89 nmol*h/mg (ND2), and 1.56 nmol*h/mg (ND3), while it was below the limit of detection in all GLD fibroblasts, indicating that the mutated GALC proteins have no residual functionality. By using an integration-free, feeder-free microfluidic reprogramming system (Luni et al., 2016) we obtained several iPSC clones from each patient and ND fibroblast line (reprogramming efficiency: 2-53%)(**Figure S1A**). After 6-10 subculturing passages, we randomly selected two clones/line for further analysis (**Figure S1A**). The presence of GALC mutations was verified by PCR analysis and sequencing (**Figure S1B-C**).

We assessed the pluripotency of GLD and ND iPSC clones by RT-PCR **(Figure S1D**) and immunocytochemistry (**Figure 1A**) for the markers NANOG, SOX2, KLF4, OCT4, and TRA1-60. Analysis of endodermal-, mesodermal-, and ectodermal-specific markers in embryoid bodies (**Figure 1B; Figure S1E**) and histological assessment of teratomas (**Figure S1F**) confirmed the ability of GLD and ND iPSCs to differentiate in multiple cell types and tissues both *in vitro* and *in vivo*.

**Figure 1.**
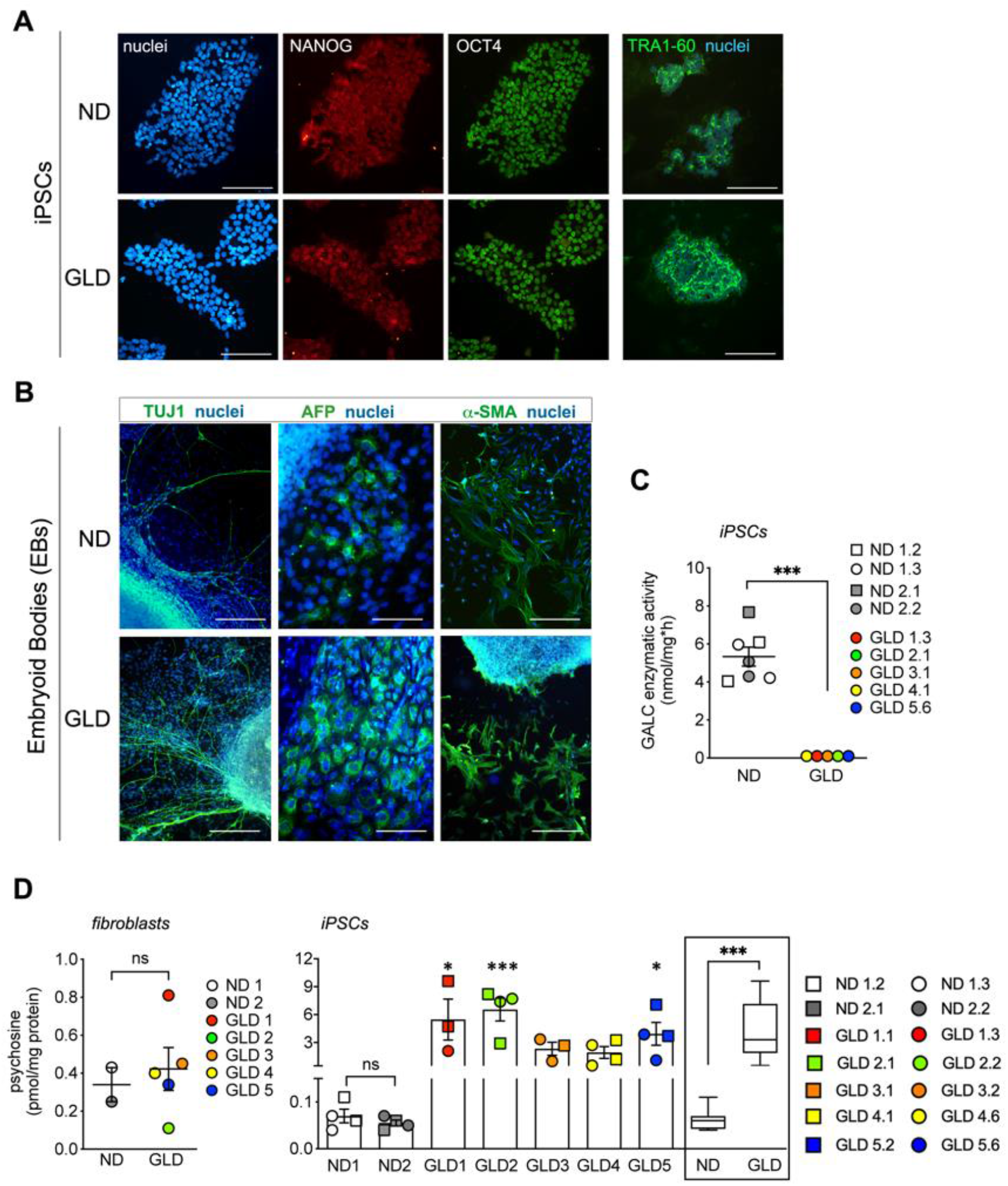
GLD iPSCs have undetectable GALC activity and increased psychosine levels (see also Figures S1 and S2) A) Representative confocal immunofluorescence pictures of ND and GLD iPSCs expressing OCT4 (green), NANOG (red), and TRA 1-60 (green). Nuclei are counterstained with Hoechst (blue). Single channels (OCT4, NANOG, nuclei) and the merged picture (TRA 1-60/nuclei) are shown. Scale bars: 100μm. B) Representative immunofluorescence pictures showing cells expressing endodermal (AFP), mesodermal (α-SMA), and ectodermal (TUJ1) markers in ND and GLD embryoid bodies (EBs). Lineage markers in green, nuclei counterstained with Hoechst (blue). Scale bars: 80 μm (AFP) and 200 μm (TUJ1 and α-SMA). C) GALC enzymatic activity in ND and GLD iPSC clones. Data are expressed as the mean ± SEM and analyzed using Mann-Whitney test; ***p<0.001. D) Psychosine levels measured by mass spectrometry in ND1 (white circles), ND2 (grey circles), and GLD (colored circles) fibroblast cell lines (n=1 sample/line) and the corresponding iPSC clones (n=2 clones/line in single or duplicate). Data are expressed as the mean ± SEM and analyzed by Mann-Whitney test (fibroblasts) and Kruskal Wallis test followed by Dunns’ multiple comparison test (iPSCs); ns, not significant * p<0.05, ***p<0.001 vs ND (mean). The mean ± SEM of all ND and GLD clones are plotted in the box and whiskers graph and analyzed by Mann-Whitney test; ***p<0.001.

All the GLD iPSC clones displayed undetectable GALC activity (**Figure 1C**) and significant psychosine storage (50- to −100-fold the ND levels) that was absent in parental fibroblasts (**Figure 1D).**

Overall, GLD patient iPSCs showed the primary biochemical GLD hallmarks while retaining normal stem cell properties, justifying their use as human-based GLD model.

### LV-mediated gene transfer in GLD iPSCs rescues GALC activity and normalizes psychosine storage

We focused on the mutant lines GLD1 and GLD5 (two clones/line) for further characterization and gene transfer studies. GLD1 is homozygous for the large 30 Kb *GALC* gene deletion (c.1161+6532_polyA+9kbdel) producing a truncated protein that lacks the whole coding region for the 30kDa subunit and 15% of the coding region for the 50kDa subunit (Luzi et al., 1995; Tappino et al., 2010). GLD5 is homozygous for a missense mutation (c.1657G>A) that is predicted to cause GALC protein misfolding (De Gasperi et al., 1996; Tappino et al., 2010). ND1 and ND2 iPSCs (two clones/lines) were used as controls for all the subsequent experiments (**Table S1, Figure S1A**).

We selected ND and GLD iPSC clones with normal karyotype (**Figure S2**) and used these cells within subculturing passages 8-30. To restore GALC activity, we transduced GLD1 and GLD5 iPSC clones using a laboratory grade/scale VSV-pseudotyped third-generation lentiviral vector (LV) encoding for the human (h) *GALC* gene tagged with the Myc peptide under the control of the human phosphoglycerate kinase (PGK) promoter (LV.hGALC) (Meneghini et al., 2016)(**Figure 2A**). Pilot dose-response experiments showed the proficient transduction of iPSC clones at Multiplicity of Infection (MOI)=2 (**Figure S3A**), which was selected to transduce GLD1.1, GLD1.3, GLD5.2, GLD5.6 and ND2.2 iPSC clones in all the following experiments. LV.hGALC-transduced GLD (GLD^GALC^) and ND (ND^GALC^) iPSCs showed vector copy number (VCN) ranging from 5 to 15 (**Figure 2B**), significant GALC mRNA (**Figure 2C**) and protein expression (**Figure 2D-E),** and increased enzymatic activity (reaching supraphysiological levels; **Figure 2F**). We observed a clear correlation between VCN, GALC mRNA expression and enzymatic activity (**Figure S2B**).

**Figure 2.**
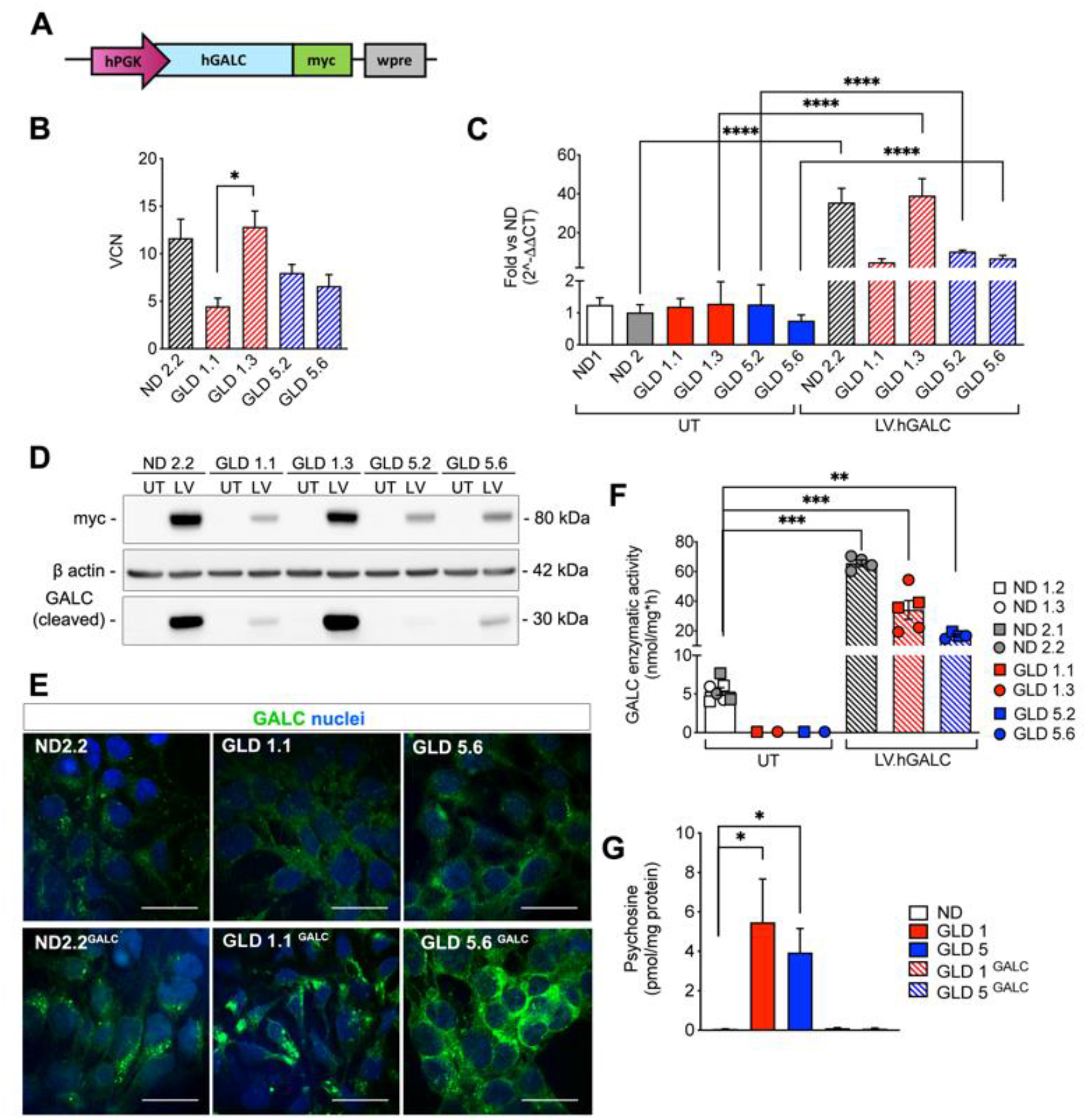
LV-mediated gene transfer rescues GALC expression and enzymatic activity in GLD iPSCs (see also Figure S3) A) Schematic representation of the LV.hGALC vector. hPGK, human phosphoglycerate kinase promoter; wpre, Woodchuck hepatitis virus Post-transcriptional Regulatory Element. B) Vector Copy Number measured in ND and GLD iPSCs clones transduced with LV.hGALC at multiplicity of infection= 2 (MOI2). Data are expressed as the mean ± SEM (n=2-4 replicates/clone) and analyzed using Kruskal Wallis test followed by Dunnett’s multiple comparison post-test; *p<0.05. C) GALC mRNA expression in untreated (UT; solid bars) and LV.hGALC-transduced (striped bars). n=3 replicates/clone. Values are normalized on GAPDH and expressed as fold change (2^-ΔΔCT) vs ND (mean). Data are expressed as mean ± SEM and analysed using one-way Anova followed by Turkey multiple comparison post-test; **** p<0.0001. D) Representative Western Blot showing the expression of the GALC-myc fusion protein using an anti-myc antibody (band size 80 kDa, precursor protein and an anti-hGALC antibody (band size 30 kDa, cleaved protein) in LV.hGALC transduced (LV; MOI2) ND and GLD iPSCs clones. β-actin was used as loading control. Note that higher protein expression in clones ND2.2 and GLD 1.3, which show higher VCN (B) and GALC activity (C, F). Untransduced (UT) clones do not express the myc-tagged protein (80kDa). The 30KDa band is undetectable in UT clones due to the low levels of physiological protein expression (see also panel E). E) Representative immunofluorescence pictures showing the expression of the GALC protein (green; anti-hGALC) in untransduced (UT) and LV.hGALC transduced ND and GLD iPSCs. Nuclei are counterstained with Hoechst (blue). Scale bars= 20μm. F-G) GALC enzymatic activity (F) and Psychosine levels (G; clones as in F) in untransduced (UT, solid bars) and LV.hGALC transduced (striped bars) ND, GLD and iPSCs. Data are expressed as the mean ± SEM (n=2-4 replicates/clone). Data were analyzed by Kruskal Wallis test followed by Dunnett’s multiple comparison post-test (ND selected as control group); **p < 0.01; ***p < 0.001; ns, not significant.

The transgenic GALC enzyme was functional and able to normalize psychosine levels in GLD iPSCs (**Figure 2G**), which do not show other pathological hallmarks related to GALC deficiency (i.e. GalCer accumulation, increased LAMP1 expression – index of lysosomal expansion) (**Figure S3C-D).** Importantly, LV transduction and GALC overexpression did not affect the pluripotency (**Figure S3E**) and proliferation rate (**Figure S3F**) of ND and GLD iPSC clones.

### GLD and ND iPSC-derived NPCs are generated with similar efficiency and show comparable phenotype

We next assessed the impact of GALC deficiency and LV-mediated GALC restoration in iPSC-derived neural cells. We first differentiated GLD, GLD^GALC^, ND, and ND^GALC^ iPSCs into NPCs taking advantage of a published protocol (Chambers et al., 2009)(**Figure 3A**). iPSC-NPCs showed downregulation of pluripotency markers (OCT4, NANOG) and upregulation of NPC markers (NESTIN, PAX6, FOXG1), early neuronal (Doublecortin, DCX) and glial cell (ASCL1) markers **(Figure 3B-C).** We obtained NPCs with similar efficiency from untransduced (UT) and LV.hGALC-transduced GLD and ND iPSCs in multiple rounds of differentiation experiments with the only exception of clone GLD1.3^GALC^, which consistently displayed abnormal morphology (not shown) and modest PAX6 upregulation as compared to other clones (**Figure 3B**). Based on these observations suggesting incomplete/defective neural induction, we excluded clone GLD1.3^GALC^ from further analysis. All the selected NPC lines stably expanded in culture for 5-6 passages in the presence of neural medium (N2)(**Figure 3D).**

**Figure 3.**
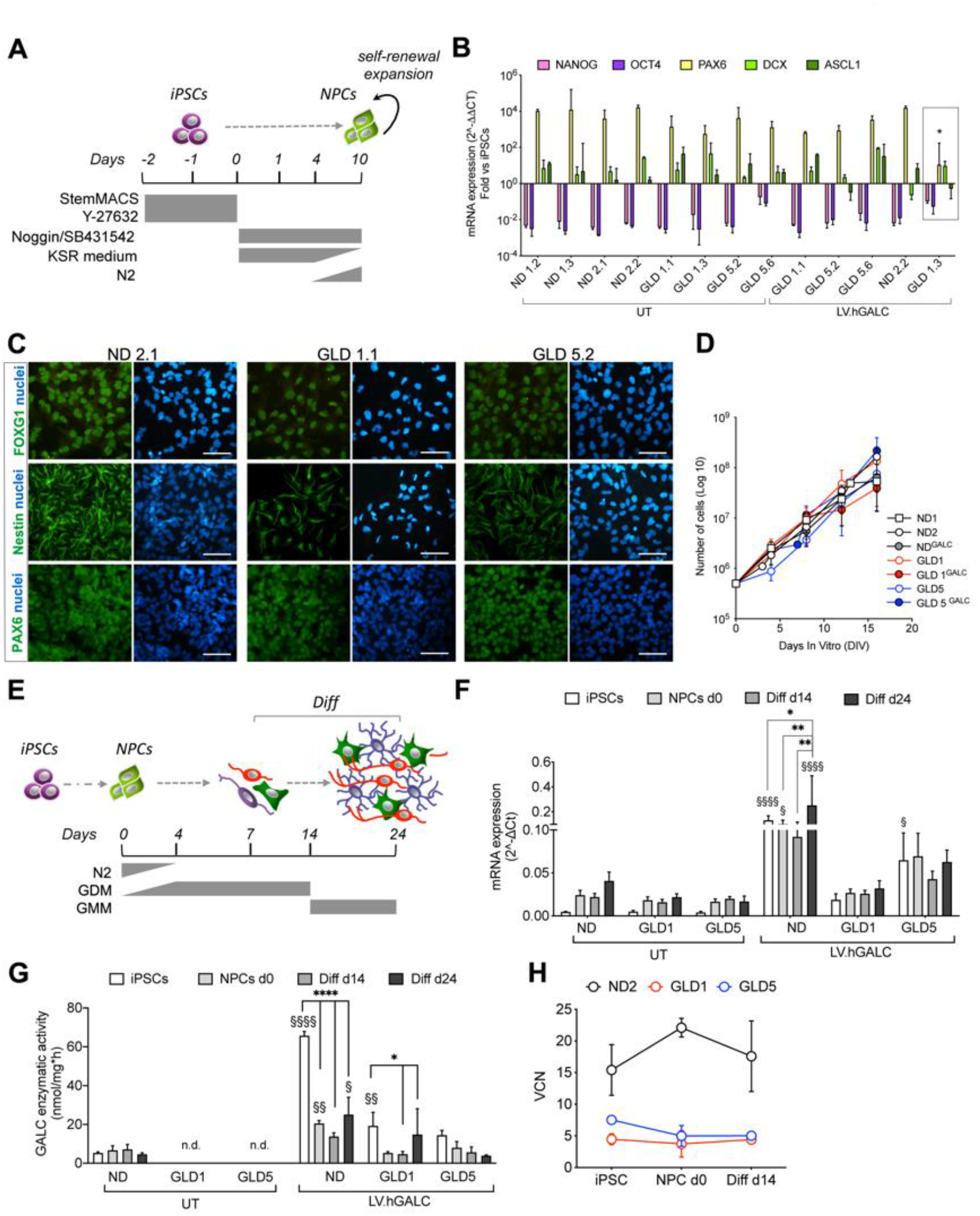
Differentiation of ND and GLD iPSCs into NPCs, neurons and glial cells. A) Schematic representation of the protocol used to differentiate iPSCs into NPCs. B) Downregulation of the expression of pluripotency-associated markers (NANOG, OCT4) and upregulation of the expression of NPC markers (PAX6, DCX, ASCL1) in untransduced (UT) and LV.hGALC-transduced ND and GLD iPSC-derived NPC populations (assessed by SYBR green RT-PCR). Data are normalized on GAPDH expression and shown as fold change (2^-ΔΔCT) on the corresponding iPSC clone. Data are expressed as the mean ± SEM (n=3-4 independent experiments; 2-3 replicates/clone). The GLD1.3 clone (box) showed reduced upregulation of PAX6. Data analyzed using Kruskal Wallis test followed by Dunnett’s multiple comparison post-test (ND selected as control group). *p<0.05. C) Representative immunofluorescence pictures showing the expression of the neural markers Nestin, FOXG1 and PAX6 (green) in ND and GLD iPSC-NPC cultures. Nuclei are counterstained with DAPI (blue). Scale bars=50μm. D) Proliferation and stable expansion rate of untransduced and LV.hGALC transduced ND and GLD iPSC-NPC lines (subculturing passages 0 to 5 shown in the graph; 16 days in vitro, DIV). Each point of the curves is the mean ± SEM; n=2-3 independent experiments, 1-2 clones/group. Clones used: ND1.2, ND1.3, ND2.1, ND2.2, GLD1.1, GLD1.3, GLD5.2, GLD5.6. E) Schematic representation of the differentiation protocol used to differentiate iPSC-NPCs (d0) into mixed neuronal/glial cultures that were analyzed in at day 7 (d7), d14, and d24. GDM, Glial Differentiation Medium; GMM, Glial Maturation Medium (see methods). F) GALC mRNA expression in untransduced (UT) and LV.hGALC-transduced iPSCs and neural progeny (NPCs, d0; differentiated cells, d14 and d24) n.d.: not detectable. Values are normalized on GAPDH and expressed as (2^-ΔCT). Clones used: ND1.2, ND1.3, ND2.1, ND2.2, GLD1.1, GLD1.3, GLD5.2, GLD5.6 Data are expressed as the mean ± SEM (n=2-3 independent experiments; 1-2 replicates/clone). Data analyzed using two-way ANOVA. To compare the different time points (iPSC, NPC d0, differentiated cells, d14 and d24) in each group we used Turkey’s multiple comparison post-test (*p < 0.05, **p < 0.01, ****p < 0.0001); to compare the different groups at each time point we used Dunnett’s multiple comparison post-test (ND selected as control group). §p < 0.05, §§p < 0.01, §§§§p < 0.0001. G) GALC enzymatic activity in untransduced (UT) and LV.hGALC-transduced ND and GLD iPSCs and neural progeny (NPCs, d0; differentiated cells, d14 and d24). Data are expressed as the mean ± SEM (n=2-4 replicates/clone). Data were analyzed by Kruskal Wallis test followed by Dunnett’s multiple comparison post-test (ND selected as control group); *p < 0.01; ****p < 0.0001; ns, not significant. H) Vector copy number (VCN) measured in LV.hGALC-transduced iPSCs and neural progeny (NPCs, d0; differentiated cells, d14). Clones used: ND2.2^GALC^, GLD1.1^GALC^, GLD5.2^GALC^, GLD5.6^GALC^; n= 1-3 independent experiments, 1-2 clones/group.

Thus, ND and GLD iPSC-derived NPCs display bona-fide NPC molecular and phenotypic features.

### Time- and cell type-dependent transcriptional regulation of GALC expression and activity during iPSC-NPC differentiation to neurons and glia

Myelinating cells and neurons are highly susceptible to psychosine-mediated toxicity, but astrocytes are also affected (Misslin et al., 2017; Potter and Petryniak, 2016). Therefore, we envisaged that differentiation of iPSC-NPCs into mixed neuronal/glial cultures would better model the composite GLD pathology. We adapted a published protocol (Frati et al., 2018) to generate cell populations containing neurons, astrocytes, and oligodendrocytes from ND, GLD, GLD^GALC^, and ND^GALC^ iPSC-NPCs. Cultures were evaluated at day 0 (d0; NPCs), d7, d14, and d24 of differentiation (**Figure 3E**), to track the time-course GALC expression/activity and the glial/neuronal cell commitment and maturation by means of biochemical and molecular analysis.

The GALC transcript was expressed at low levels in both ND and GLD iPSC clones (GLD1 and GLD5 mutations are not predicted to impact on mRNA transcription) and was upregulated during the neural induction and neuronal/glial differentiation (**Figure 3F**). GALC mRNA expression in GLD^GALC^ and ND^GALC^ iPSCs was constant over time, due to the high and stable expression driven by the constitutive PGK promoter **(Figure 3F).**

ND NPCs and differentiated cultures (d14 and d24) showed comparable GALC enzymatic activity **(Figure 3G).** The supraphysiological (≈10X the normal levels) GALC activity in GLD^GALC^ iPSCs progressively decreased during the neural differentiation to reach physiological levels at d14 and d24, whereas ND^GALC^ differentiated cultures maintained supraphysiological GALC activity (≈3-fold the ND counterpart at d14 and d24) (**Figure 3G**). The stable VCN measured in ND^GALC^ and GLD^GALC^ cultures at different time points of differentiation (**Figure 3H**) rules out the possibility that the decrease in enzymatic activity results from the counter selection of cells harboring high VCN.

Overall these data show that LV-mediated gene transfer ensures (supra)physiological GALC expression and activity in iPSC-NPCs and neuronal/glial progeny, and highlight a stringent transcriptional regulation of GALC activity during the iPSC to neural differentiation.

### GLD iPSC-NPCs display defective neural and glial differentiation: impact of GALC rescue/overexpression on psychosine levels and phenotype

The altered expression pattern of the master oligodendroglial gene OLIG2 observed in GLD1 and GLD5 cells as compared to ND cells suggest an altered oligodendroglial differentiation program, which was only partially normalized in GLD^GALC^ cultures **(Figure S4A)**. Interestingly, a similar (or even more severe) alteration was observed in ND^GALC^ cultures, in which OLIG2 expression is overall strongly reduced (**Figure S4A**). Immunofluorescence and WB analysis showed an early and persistent reduction of oligodendroglia (OLIG2+, APC+ cells; **Figure 4A-B)** and a significant downregulation of OLIG1 protein expression (**Figure S4B**; d24) in both GLD1 and GLD5 NPC-derived cultures as compared to the ND counterpart, highlighting a partial phenotypic rescue in GLD^GALC^ cells and a detrimental impact of GALC overexpression in ND cultures (**Figure 4A-B** and **Figure S4B).**

**Figure 4.**
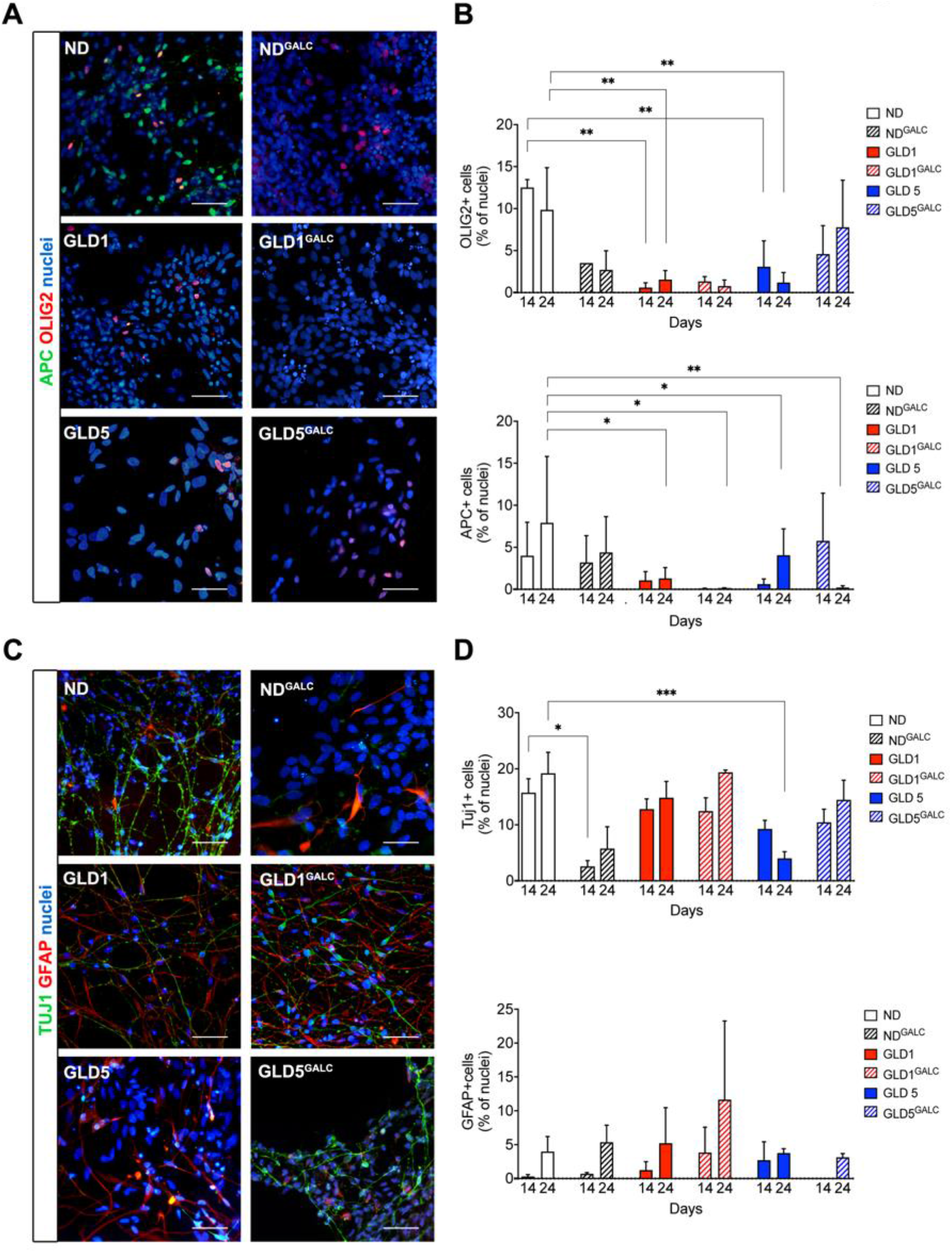
GLD NPCs display impaired neuronal and glial differentiation (see also Figures S4, S5, S6, S7) A) Representative pictures of untransduced (UT) and LV.hGALC-transduced ND and GLD differentiated neural progeny (d24 of differentiation) expressing Olig2 (red) and APC (green). Nuclei are counterstained with Hoechst (blue). Scale bar, 50 μm. B) Quantification of cells expressing Olig2 and APC in UT and LV.hGALC-transduced ND and GLD differentiated neural progeny (d14 and d24). C) Representative pictures of UT and LV.hGALC-transduced ND and GLD differentiated neural progeny (d24 of differentiation) expressing TUJ1 (green; neurons) and GFAP (red; astrocytes). Nuclei are counterstained with Hoechst (blue). Scale bar, 50 μm. D) Quantification of cells expressing TUJ1 and GFAP in UT and LV.hGALC-transduced ND and GLD differentiated neural progeny (d14 and d24). Data in B and D are expressed as percentage of immunoreactive (IR) cells on total nuclei and represent the mean+SEM; n= 2-4 independent experiments; 1-3 clones/group with 2-3 replicates/clone for each time point; 5-10 fields/coverslips were analyzed. Clones used: ND1.2, ND2.1, ND2.2, GLD1.1, GLD1.3, GLD5.2, GLD5.6, ND2.2^GALC^, GLD1.1^GALC^, GLD5.2^GALC^, GLD5.6^GALC^. Data were analyzed by two-way ANOVA followed by Dunnett’s multiple comparison post-test (ND at the corresponding time point selected as control group). *p<0.05, **p<0.01, *** p < 0.001.

GALC deficiency affected neuronal differentiation, with some differences observed in the two GLD lines. The mRNA expression of Doublecortin (DCX), a marker of immature neurons, was reduced in GLD5 (but not in GLD1) as compared to ND cells **(Figure S5A).** In line with these results, GLD5 cultures showed a significant time-dependent reduction of TUJ1+ neurons (**Figure 4C-D**) and a decreased and/or delayed TUJ1 and neurofilament (NF)-M protein expression, respectively (**Figure S5B-C).** The major decrease in the expression of acetylated and polyglutamated tubulin and phosphorylated NF observed in GLD5 cultures (d14 and d24; **Figure S6**) further suggested neuronal loss and/or defective neuronal differentiation/maturation. In contrast, GLD1 cultures displayed normal proportions of neuronal cells (**Figure 4C-D)** and normal expression of neuronal proteins (**Figure S5B-C)**. The quantitative and qualitative neuronal defects observed in GLD5 cultures were rescued in differentiated GLD^GALC^ cultures (**Figure 4C, Figure S5B-C**), which also showed an early upregulation of NF-M protein as compared to the ND cultures and untransduced counterparts (**Figure S5C),** suggesting an accelerated neuronal differentiation associated to GALC overexpression. Similarly to what observed in oligodendroglial cells, GALC overexpression negatively affected the pattern of gene and protein expression (**Figure S5, S6**), as well as the number and morphology of ND neurons (**Figure 4C-D)**.

Astroglia is the less represented and most variable cell population in both ND and GLD differentiated cultures, with no clear differences related to GALC deficiency, as shown by glial fibrillary acidic protein (GFAP) gene expression analysis (**Figure S7A**), WB analysis (**Figure S7B**), and IF **(Figure 4C-D** and **Figure S7C**; 3-15% of GFAP+ cells on the total cells at d24).

Interestingly, both GALC deficiency (GLD cultures) and forced GALC expression (GLD^GALC^ and ND^GALC^ cultures) were associated to higher percentages of cells that stained negative for the selected lineage-specific markers **(Figure S7C).**

Psychosine levels were low in GLD iPSC-NPCs and increased during differentiation, to reach ≈8 (GLD1) and ≈30 fold (GLD5) the normal levels at d24 (**Figure 5A**). The psychosine build up was not associated to a significant increase of apoptotic cells (cleaved caspase 3+; **Figure 5B**) or to enhanced phospholipase A2 (PLA2) activity (**Figure 5C**), which has been associated to psychosine-induced apoptotic cell death (Misslin et al., 2017). Psychosine levels were normalized in GLD1^GALC^ cultures and strongly reduced in GLD5^GALC^ cultures (≈2-fold the normal), in line with the enzymatic reconstitution. We observed moderate enlargement of lysosomal vesicles (Lysotracker assay) (**Figure 5D**) in GLD1 NPCs, which was normalized in GLD1^GALC^ cultures. LAMP1 protein expression was comparable in ND, GLD, and GLD^GALC^ differentiated cultures, while it was increased in GALC-overexpressing ND cells (**Figure 5E**).

**Figure 5.**
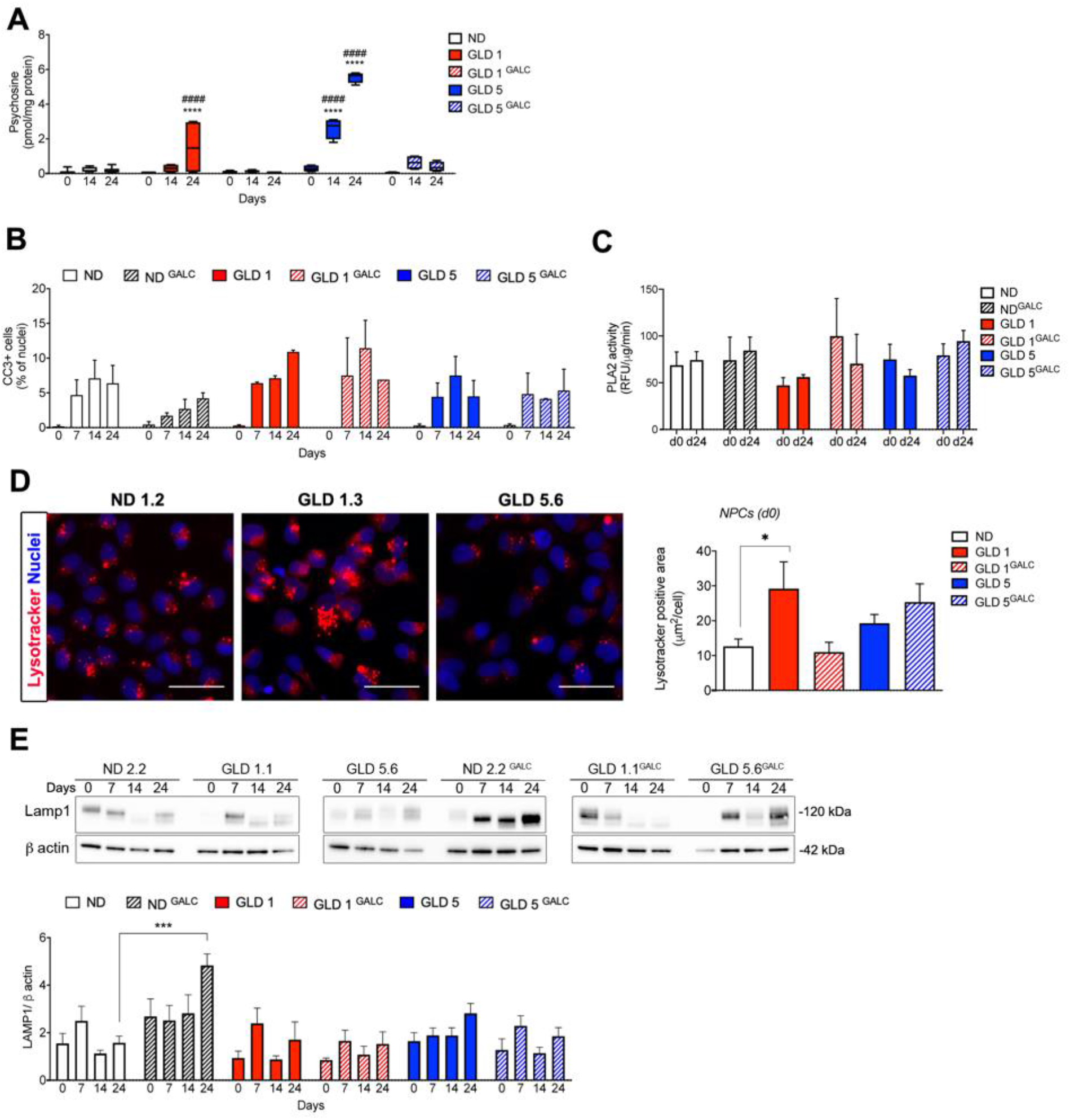
Pathological hallmarks in GLD neural progeny. A) Psychosine levels in UT and LV.hGALC-transduced ND and GLD NPCs (d0) and differentiated progeny (d14 and d24). Data are expressed as mean+SEM, n=2-4 replicates/clone, 1-3 clones/group. Clones used: ND1.2, ND2.1, ND2.2, GLD1.1, GLD1.3, GLD5.2, GLD5.6, ND2.2^GALC^, GLD1.1^GALC^, GLD5.2^GALC^, GLD5.6^GALC^. Data were analyzed by two-way ANOVA followed by Turkey’s multiple comparison post-test; ****p<0.0001 GLD vs ND at the corresponding time point; #### p<0.0001 GLD vs GLDGALC at the corresponding time point. p>0.05 GLD^GALC^ vs ND at all time points. B) Quantification of cells expressing cleaved caspase 3 (CC3) in UT and LV.hGALC-transduced ND and GLD NPCs (d0) and differentiated progeny (d14 and d24). Data are expressed as percentage of immunoreactive (IR) cells on total nuclei and represent the mean+SEM; n= 2 independent experiments; 1-3 clones/group, 2-3 replicates/clone for each time point; 5-10 fields/coverslips were analyzed. Clones used: ND1.2, ND2.1, ND2.2, GLD1.1, GLD1.3, GLD5.2, GLD5.6, ND2.2^GALC^, GLD1.1^GALC^, GLD5.2^GALC^, GLD5.6^GALC^. Data were analyzed by two-way ANOVA followed by Dunnett’s multiple comparison post-test (ND at the corresponding time point selected as control group). No differences between groups (p>0.05). C) PLA2 activity in untransduced and LV.hGALC-transduced ND and GLD NPCs (d0) and differentiated progeny (d24). Data are expressed as mean+SEM. n=2 independent experiments; 1-3 clones/group. Clones used: ND1.2, ND2.1, ND2.2, GLD1.1, GLD1.3, GLD5.2, GLD5.6, ND2.2^GALC^, GLD1.1^GALC^, GLD5.2^GALC^, GLD5.6^GALC^. D) Representative pictures of lysotracker-positive NPCs (red) and quantification of lysotracker-positive area (μm^2^/cell) in ND1.2, GLD1.3 and GLD5.6 clones. Nuclei are counterstained with Hoechst. Scale bars= 20μm. Data are expressed as the mean+SEM, n=2 independent experiments, 3 wells/clone, 1.400-12.000 cells/well were analyzed. Data analysed by two-way ANOVA followed by Dunnett’s multiple comparison post-test (ND selected as control group). *p<0,05. E) Representative Western Blot and densitometric quantification showing the expression of the LAMP1 protein in UT and LV.hGALC-transduced ND and GLD NPCs (d0) and differentiated progeny (d7, d14 and d24). β-actin was used as loading control. Data are represented as the mean+SEM; n= 3-5 independent experiments; 1-4 clones/group. Data were analyzed by two-way ANOVA followed by Dunnett’s multiple comparison post-test (ND at the corresponding time point selected as control group). * p<0.05, **p< 0.01; *** p < 0.001.

Taken together, these data show that: i) GALC deficiency severely impairs the differentiation of hiPSC-derived NPCs into oligodendrocytes and neurons, with cell type- and patient-specific phenotypes; ii) LV-mediated GALC reconstitution in GLD cells reduces psychosine levels and provides partial phenotypic rescue in GLD neural progeny; iii) GALC-overexpressing ND cells display lysosomal expansion and a defective neuronal and glial phenotype, suggesting that a tight regulation of GALC expression is required to ensure proper NPC functionality and preserve multipotency.

### Early activation of a senescence response in GLD5 iPSC-derived neural cells

Our data suggested a marginal contribution of cell death to the loss of GLD neurons and glial cells. Therefore, we explored whether the altered differentiation programs of GLD NPCs and progeny could be caused by an alteration of proliferative potential. By performing and quantifying Ki67/nestin immunostaining at the different stages of the differentiation protocol, we showed a time-dependent decrease in the percentage of proliferating cells in all ND and GLD cultures, indicating the lineage commitment of iPSC-NPCs, their exit of cell cycle and acquisition of a post-mitotic state (**Figure 6A).** Interestingly, EdU incorporation assay performed at the same time points during differentiation showed an earlier and more drastic drop in the percentage of cells in active S-phase in GLD5 cultures as compared to ND and GLD1 counterparts (**Figure 6B).** This proliferation drop was recovered in GLD5^GALC^ cultures (**Figure 6B**).

**Figure 6.**
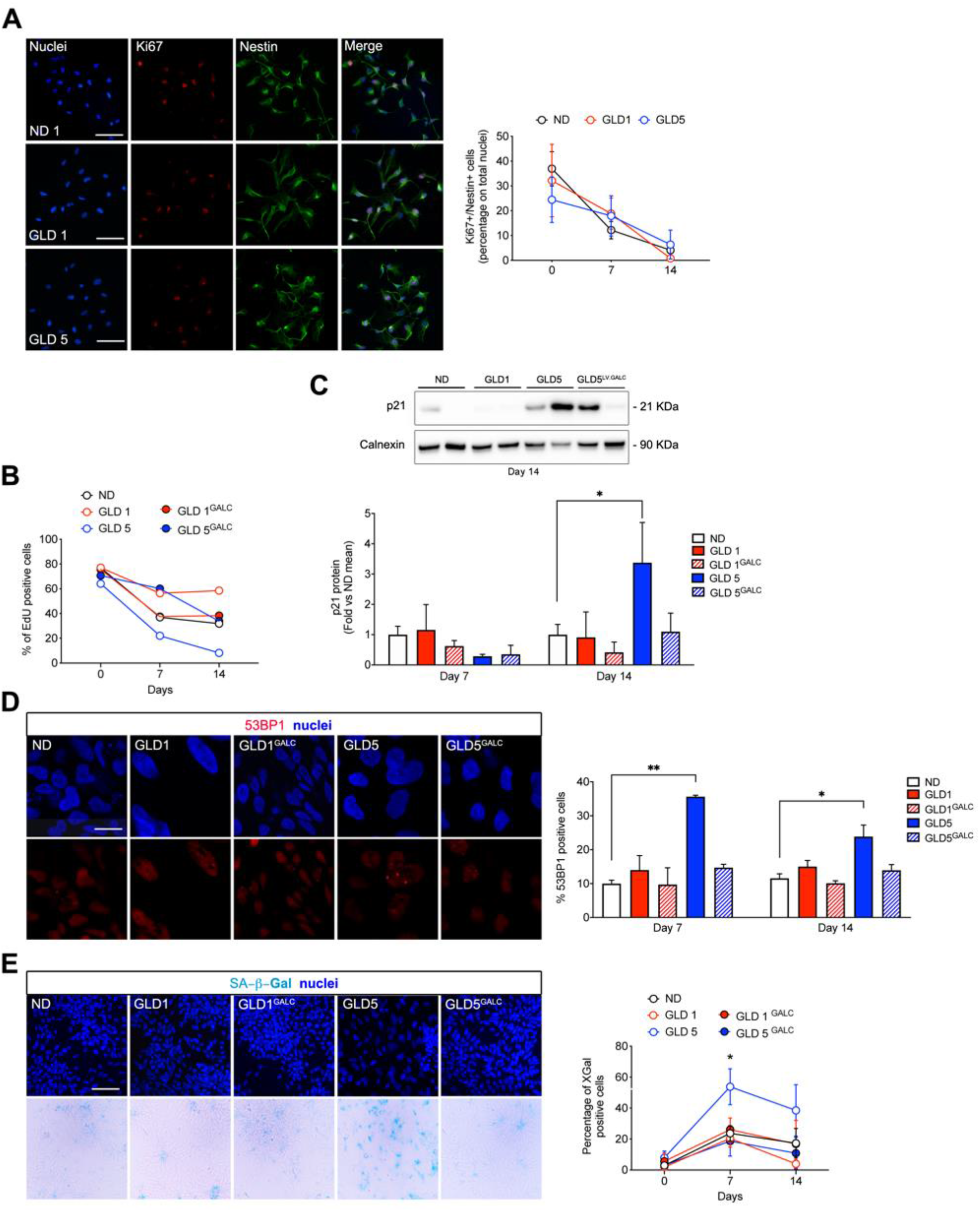
GLD5 NPCs progeny display markers of senescence (see also Figure S8) A) Representative confocal immunofluorescence pictures and quantification of cells expressing nestin (green) and Ki67 (red) in ND, GLD, and GLD^GALC^ NPCs (d0; shown in the pictures) and differentiated cultures (d7 and d14). Nuclei are counterstained with Hoechst. Scale bars= 50μm. Data in the graph are expressed as the mean+SEM n= 3 independent experiments; 1-3 clones/group with 1-3 replicates/clone for each time point; 5-10 fields/coverslips were analyzed. Clones used: ND1.2, ND1.3, ND2.1, ND2.2, GLD1.1, GLD1.3, GLD5.2, GLD5.6). B) Quantification of EdU positive cells (after a 16 hour-pulse) in ND, GLD, and GLD^GALC^ NPCs (d0) and differentiated cultures (d7 and d14). Data in the graph are expressed as percentage over total cells (mean); n= 2 independent experiments for each time point, 1-2 clones/group. Clones used: ND1.2, ND2.2, GLD1.1, GLD5.2, GLD5.6, GLD1.1^GALC^, GLD5.2^GALC^. C) Representative Western blot and densitometric quantification showing the expression of the p21 protein in ND, GLD, and GLD^GALC^ NPC differentiated progeny (d7 and d14; d14 shown in the blot). Calnexin was used as loading control. Data are expressed as fold vs ND at the corresponding time point (mean of all clones) and represented as the mean+SEM; n= 2 independent experiments; 1-3 clones/group. Clones used: ND1.2, ND2.2, GLD1.1, GLD1.3, GLD5.2, GLD5.6, GLD1.1^GALC^, GLD5.2^GALC^, GLD5.6^GALC^. Data were analyzed by two-way ANOVA followed by Dunnett’s multiple comparison post-test (ND at the corresponding time point selected as control group). * p<0.05. D) Representative confocal immunofluorescence pictures and quantification for 53BP1-positive DDR foci in ND, GLD, and GLD^GALC^ NPC (d0) and differentiated cultures (d7 and d14; d14 shown in the pictures). Nuclei are counterstained with DAPI. Scale Bar = 15 μm (for all panels, shown in ND). Clones used: ND1.2, ND2.2, GLD1.1, GLD1.3, GLD5.2, GLD5.6, GLD1.1^GALC^, GLD5.2^GALC^, GLD5.6^GALC^. Data in the graph are expressed as the mean+SEM; n= 3 independent experiments, 2-3 replicates/clone for each time point. 100 to 600 nuclei per condition were analyzed. Data were analyzed by Kruskal Wallis test followed by Dunns’ multiple comparison post-test (ND at the corresponding time point selected as control group). * p<0.05, **p < 0.01. E) Representative pictures (brightfield and immunofluorescence for nuclei) and quantification of SA-βGal-positive cells in ND, GLD, and GLD^GALC^ NPCs (d0) and differentiated cultures (d7 and d14; d7 shown in the pictures). Nuclei are counterstained with Hoechst. Scale bars= 50μm (for all panels, shown in ND). Data in the graph are expressed as the mean+SEM n= 3 independent experiments; 1-3 clones/group, 1-3 replicates/clone for each time point; 4-10 fields/coverslips were analyzed. Clones used: ND1.2, ND2.1, ND2.2, GLD1.1, GLD5.2, GLD5.6, GLD1.1^GALC^, GLD5.2^GALC^, GLD5.6^GALC^. Data were analyzed by two-way ANOVA followed by Dunnett’s multiple comparison post-test (ND at the corresponding time point selected as control group). * p<0.05.

We next investigated whether the early proliferation arrest highlighted in GLD5 cultures was associated with the establishment of cellular senescence, a biological process characterized by stable growth arrest, accumulation of DNA damage and activation of a pro-inflammatory program collectively known as SASP, senescence-associated secretory phenotype (He and Sharpless, 2017). We detected upregulation of the cell cycle inhibitor p21 protein (**Figure 6C)** and increased nuclear levels of DNA damage response (DDR) markers 53BP1 (**Figure 6D)** and yH2AX **(Figure S8A-B)** as early as d7 post-differentiation. The induction of DDR and p21 was specific of GLD5 cultures, not reported in ND or GLD1 cultures, and was partially rescued by in GLD5^GALC^ cultures. We also observed a dramatic increase in the percentage of Senescence Associated (SA) β-gal positive cells in GLD5 (but not ND or GLD1 cultures) with a peak at d7 of differentiation (**Figure 6E**), and variable induction of the SASP pro-inflammatory factors TNF-α and IL1-β (**Figure S8E-F**). These senescence-associated hallmarks were reduced in GLD5^GALC^ cultures.

Altogether, our data suggest that GLD5 iPSC-derived NPCs activate an early senescence program that hampers physiological lineage commitment/differentiation. This senescent phenotype is patient-specific (as it is not evident in GLD1 cells), and is rescued by supplying physiological levels of a functional GALC enzyme.

### GLD cells display an altered lipidomic profile during the iPSC to neural differentiation

Sphingolipids display widespread developmental, physiological and pathological actions in the CNS, and induce biological effects on various stem/progenitor cell types (Hannun and Obeid, 2017). Given the central role of GALC in the sphingolipid metabolism, we performed an untargeted global lipidomic profiling in ND, GLD, and GLD^GALC^ iPSCs, NPCs and neuronal/glial cell progeny (d14 of differentiation) to elucidate the impact of GALC absence or enhanced expression during the iPSCs to neural differentiation. Lipid extracts were normalized based on protein concentration and analyzed by LC-MS-based lipidomics workflow. We determined the lipid species showing reproducible changes in two independent profiling experiments with two biological replicates for each experiment.

We identified more than 800 lipid species, with a distinct signature and enrichment of specific lipid classes in iPSCs (e.g. triacylglycerols, TAG; lyso-phosphatidylserine, LPS), NPCs (e.g. phosphatidylcholine, PC), and differentiated progeny (e.g phosphatidylethanolamine, PE; sphingomyelin, SM) (**Figure 7A**). We observed profound changes in a number of phospholipid classes in GLD as compared to ND cells (i.e. TAG; Diacylglycerols, DAG; Monoacylglycerols, MAG; PC; PE; Lysophosphatidylethanolamine, LPE; PS; SM; Ceramides, Cer; HexosylCeramides, HexCer)(**Figure 7B**). Within the same lipid class, we detected differences related to the stage of differentiation (iPSCs vs. NPCs. vs differentiated cells) and genotype (ND vs. GLD1 vs. GLD5). GLD1 and ND iPSCs often showed a comparable abundance of all these lipid classes, while GLD5 iPSCs displayed a clearly different trend (**Figure 7B**). The most striking differences between GLD1 and ND cells were observed in NPCs, which showed decreased levels of PE/LPE and increased Cer and DAG levels. The different lipidomic profile displayed by GLD1 and GLD5 cells become more evident as cells undergo differentiation. Indeed, when compared to GLD1 and ND cells, GLD5 cells showed reduced PC, PE, HexCer, and SM, and a strong accumulation of DAG, TAG, Cer, and PS **(Figure 7B**). The inverse pattern of Cer and SM expression in GLD5 cell populations - and particularly in differentiated cells - suggests that excess Cer could result from the SM-based synthetic pathway in these cells (Young et al., 2012).

**Figure 7.**
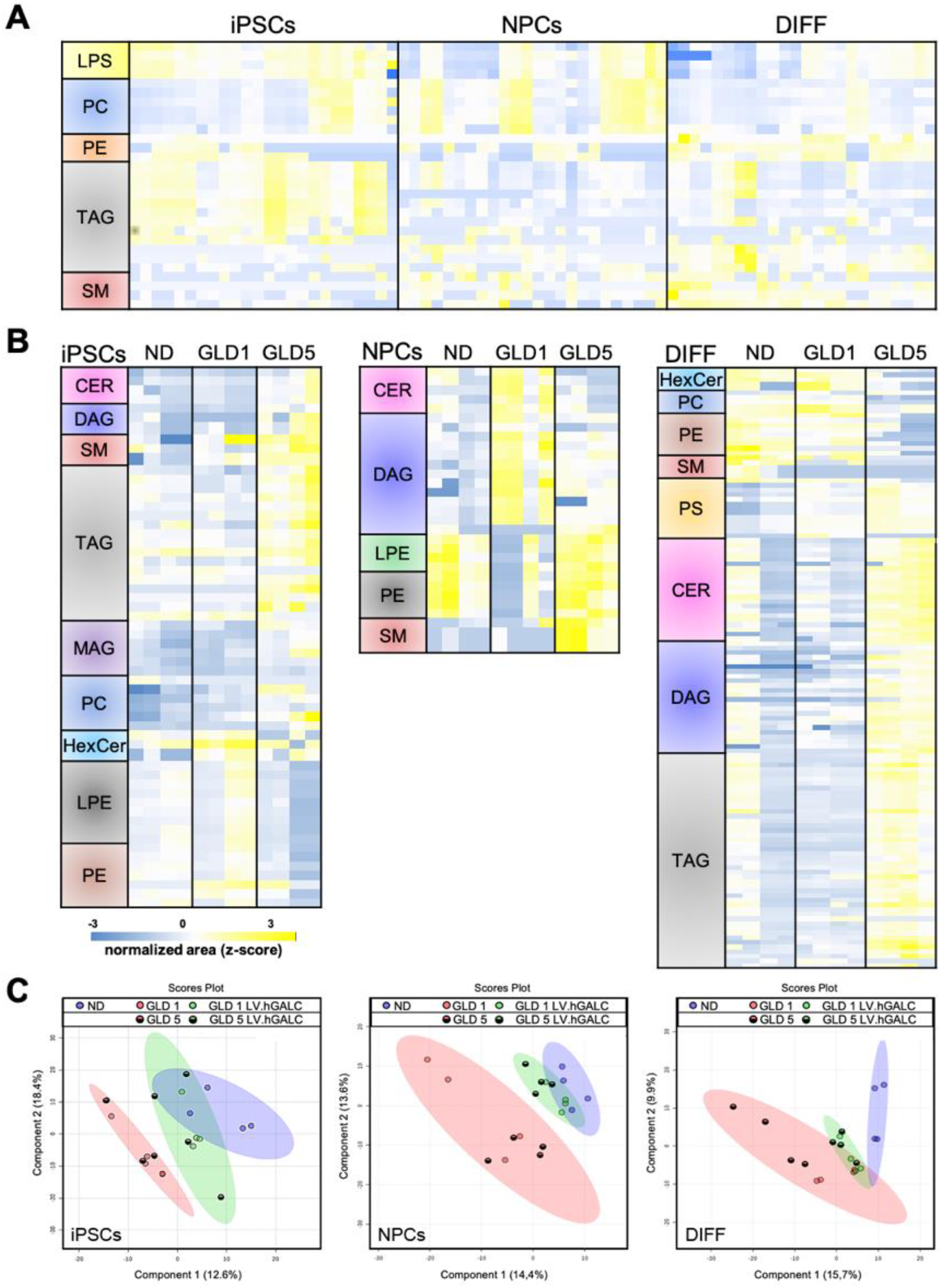
Lipidomic profile during iPSC neural differentiation (see also Figure S9) A) Hierarchical clustering of lipid species differently enriched in iPSCs, NPCs and differentiated cultures (d14). ANOVA statistical test was applied (FDR<0.05). B) Hierarchical clustering of lipid species differently enriched in ND, GLD1 and GLD5 iPSCs, NPCs and differentiated cultures (d14). ANOVA statistical test was applied (FDR<0.05). C) Principal component analysis (PCA) of lipid species distribution in ND, GLD and LV.hGALC transduced GLD iPSCs, NPCs and differentiated cultures (d14 of differentiation). Clones used: ND2.2, GLD1.1, GLD5.2, ND2.2^GALC^, GLD1.1^GALC^, GLD5.2^GALC^. See methods for abbreviation of lipid classes.

LV-mediated rescue of GALC activity partially normalized the lipidomic profiles of GLD1 and GLD5, as shown by the PCA analysis (**Figure 7C)** and lipid signature **(Figure S9A-B).** In GLD1^GALC^ cells we observed a rescue of PC, PE, HexCer (iPSCs) and TAG (NPCs). In GLD5 ^GALC^ cells we observed the normalization of HexCer and SM level, as well as a rescue to normal levels of all DAG, TAG and PC that were mostly affected in GLD5 differentiated cells. LV-mediated GALC overexpression resulted in a profound and global impact on the lipidome of ND cells. Coherently with the presence of supranormal GALC activity, we detected in ND^GALC^ iPSCs, NPCs, and differentiated cells decreased levels of HexCer and variable increase of Cer, but also of DAG and PS, in comparison to ND counterparts (**Figure S9C**).

## Discussion

Human iPSCs have been differentiated into neurons and/or glial cells to model the neural defects associated to several lysosomal storage diseases (LSDs) including mucopolysaccharidosis (Bayó-Puxan et al., 2018; Canals et al., 2015; Kobolák et al., 2019; Lemonnier et al., 2011; Vallejo-Diez et al., 2018), Niemann−Pick type C (Maetzel et al., 2014; Ordoñez and Steele, 2016; Trilck et al., 2013; Yu et al., 2014), Pompe disease (Higuchi et al., 2014), Gaucher disease (Panicker et al., 2012; Sun et al., 2015; Tiscornia et al., 2013), GM1 and GM2 gangliosidosis (Son et al., 2015)(Allende et al., 2018), NCL (Lojewski et al., 2014), and MLD (Frati et al., 2018; Meneghini et al., 2017). To our knowledge this is the first report describing a human iPSC-based neural model of GLD.

The five GLD patient-derived fibroblast cell lines that we used for somatic cell reprogramming have different mutations in the *GALC* gene, all of them associated to the infantile form of the disease. All the selected GLD iPSC lines displayed normal phenotypic and functional properties despite having undetectable GALC enzymatic activity and significant psychosine storage. The dispensability of lysosomal enzymatic activity for somatic cell reprogramming and iPSC maintenance as well as the tolerability to lipid storage has been described in human iPSCs in other LSD models (Canals et al., 2015; Lojewski et al., 2014; Meneghini et al., 2017; Trilck et al., 2013). The LV-mediated supply of physiological or slightly supraphysiological GALC activity (3-4 fold the ND levels) in GLD iPSCs normalizes psychosine levels, confirming the potential of gene transfer to correct the primary storage in GALC-deficient human cells. In line with our previous study (Meneghini et al., 2017), we did not experience LV-mediated toxicity in hiPSCs, which were viable and functional for several passages after transduction, maintaining pluripotency and stable GALC activity.

The GALC enzyme contributes to preserve NPCs functionality during post-natal murine CNS development (Santambrogio et al., 2012), but this aspect was never investigated in human NPCs, neurons, and glial cells. GLD iPSC-derived NPCs had levels of psychosine similar to their ND counterparts, differently from the parental GLD fibroblasts and iPSCs, in which psychosine is elevated. Interestingly, the negligible psychosine storage in human NPCs distinguishes them from their murine counterpart (Santambrogio et al., 2012). The differentiation of GLD iPSC-NPCs into neurons and glia exacerbates psychosine accumulation and highlights a pathological phenotype, in line with the relevant role of GALC during CNS differentiation (Graziano and Cardile, 2015). Specifically, we observed an early and persistent loss of oligodendroglial cells in both GLD1 and GLD5 NPC-derived cultures. GLD5 (but not GLD1) cultures also displayed a dramatic loss of neurons. A modest psychosine storage may affect murine neuronal and glial differentiation from NPCs (Castelvetri et al., 2011; Santambrogio et al., 2012; Teixeira et al., 2014) before overt toxicity mediated by storage overload (Hawkins-Salsbury et al., 2013; White et al., 2011)(Bongarzone et al., 2016; Ricca and Gritti, 2016). Interestingly, psychosine storage has been detected in the brain of GLD human foetuses (Igisu and Suzuki, 1984) and elevated psychosine blood concentration is considered a specific marker in infantile GLD patients (Escolar et al., 2017), who may display early white matter changes and functional disability (Gupta et al., 2015). The aggravated defects in neuronal/glial differentiation from human NPCs described here are in line with these clinical observations. Also, several findings in this study suggest a composite and mutation-dependent effect associated to GALC deficiency which impacts on human NPC commitment/differentiation besides psychosine toxicity: i) psychosine accumulation starts at early time points of NPC differentiation and increases as cells differentiate into neurons and glia; still, this progressive load is not accompanied by a dramatic increase of apoptotic cells; ii) higher percentages of marker-negative cells are constantly found in GLD differentiated cultures, together with altered expression of key genes involved in oligodendroglial, neuronal, and astroglial specification/maintenance, suggesting a block in cell commitment or differentiation; iii) we observe only a partial rescue of the differentiation program in LV.hGALC-transduced GLD neurons/glia - in which psychosine storage is cleared - with an evident amelioration in the neuronal compartment but far less improvement in the oligodendroglial population.

In the attempt of elucidating the link between GALC deficiency and defective neural differentiation, the mechanisms of disease correction upon gene transfer, and the mechanisms underlying the mutation-specific phenotype, we pointed our attention on the lipidome. Intriguingly, many sphingolipids involved in the GALC pathway, such as ceramide, sphingosine, and sphingosine-1-phosphate, contribute to proliferation, migration, cell fate, apoptosis (Hannun and Obeid, 2008, 2017), senescence and autophagy (Trayssac et al., 2018; Young and Wang, 2018). We previously documented the unbalance of some bioactive lipids in brain tissues of GLD mice (Santambrogio et al., 2012). Here, we show a distinctive global lipid profile in iPSCs vs. NPCs. vs. NPC progeny, regardless of the genotype (ND, GLD) or mutation (GLD1, GLD5), providing for the first time a snapshot of the changes in lipid metabolism during the iPSC to neural differentiation. By comparing the global lipidomic profile in each cell population, we observed that GLD and ND cells differ for the abundance of various fatty acids and phospolipid classes. The mild unbalance of lipid composition in GLD NPCs becomes more evident as cells differentiate. Intriguingly, GLD1 and GLD5 cell populations displayed remarkable differences in their lipidomic profile. Specifically, GLD5 iPSCs and NPC progeny show an accumulation of TAG, DAG, and ceramide, whose role in cell fate, cellular stress, ageing and replicative senescence has been proposed in multiple tissues and cell types (Hannun and Obeid, 2002, 2017; Lizardo et al., 2017, 2018; Venable et al., 1995). We envisage that this peculiar lipid perturbation contributes to the early senescence response and differentiation defects observed in GLD5 iPSC-derived neural cells. Considering that ceramide is the product of GalCer degradation by GALC, its accumulation in GLD5 lines is counterintuitive. We speculate that other major pathways leading to ceramide production may be induced in GLD5 cells, i.e. activation of sphingomyelinases and/or ceramide synthases (Claus et al., 2009; Mullen et al., 2012). The SM-ceramide pathway is particularly intriguing, since it may induce apoptosis, differentiation, and growth arrest (Jana et al., 2009). SM might be converted into ceramide via sphingomyelinase to compensate GALC deficiency. Moreover the parallel accumulation of DAG and TAG might be explained by an inhibition of the Kennedy pathway that is essential for *de novo* glycerophospholipid biosynthesis and the SM cycle in sphingolipid metabolism (Bieberich, 2012). Importantly, our data suggest that normalization of GALC activity rescues or at least ameliorates the global lipid composition of mutant cells.

The association between lysosomal dysfunction, lipid unbalance, and senescence in NPCs, neurons, and glial cells and its contribution to neurodegeneration and demyelination is emerging as an area of active scientific investigations. Specifically, by taking advantage of mouse models or post-mortem human brain samples it was reported that senescence functionally contributes to multiple sclerosis pathogenesis (Nicaise et al., 2019) as well as to cognitive defects in Alzheimer disease (Zhang et al., 2019). Notably, our study highlights the importance of performing this investigation in relevant human iPSC-based neural models, which recapitulate the patient-specific pathological hallmarks more faithfully as compared to murine models or non-neural cells.

Our study raises a number of relevant mechanistic questions. First, the respective role and potential interaction between psychosine load and lipidome perturbation in determining the different phenotype in GLD5 vs. GLD1 are not fully elucidated. One option would be the detrimental action exerted by psychosine and ceramide (and possibly other lipid classes, e.g. TAG) preferentially on oligodendrocyte and neuronal cells, respectively (Jana et al., 2009). Second, while our results suggest an association between ceramide/TAG/DAG accumulation and senescence, the mechanistic link between the GLD5 mutation and the occurrence of this specific lipid unbalance is not clear yet. Possibilities include: i) a direct effect of the mutated GALC protein on the lipidome (the investigation of the lipid-protein interactions and definition of the molecular mechanisms of pathway and enzyme regulation are rapidly expanding in the field of bioactive lipids) (Hannun and Obeid, 2017); ii) an indirect effect of another GLD-associated pathology (i.e. oxidative stress, mitochondrial dysfunction) (Ricca and Gritti, 2016)(Ballabio and Gieselmann, 2009). The use of isogenic pairs might clarify whether the GLD5 mutated protein has a dominant negative effect that contribute to the pathogenic phenotype in addition to the enzymatic deficiency.

In summary, our data show that GALC deficiency severely affects the differentiation of hiPSC-derived neural progenitors into neurons and oligodendrocytes. We highlight cell type-, and patient-specific disease hallmarks in GLD cells, including galactosylsphingolipid storage, lipidome perturbation, and activation of a senescence program. Finally, we show that a timely regulated GALC expression is critical for proper neural commitment and differentiation. We anticipate these findings to be starting point for future functional experiments aimed at further dissecting the complex and mutation-specific GLD pathology. This knowledge will pave the way to tailored gene/cell therapy strategies to enhance the correction of CNS pathology in GLD.

## Acknowledgments

We are grateful to Lucia Sergi Sergi for LV preparation and titration, Andrea Ditadi for providing the ESC line H9, Wim Kulik for psychosine analyses, Janet Elizabeth Deane for providing the anti-hGALC antibody, all the members of the Gritti’s lab for generous support and helpful discussion.

Part of this work was carried out in ALEMBIC, an advanced microscopy laboratory established by IRCCS Ospedale San Raffaele and Università Vita-Salute San Raffaele.

We thank the “Diagnosi PrePostnatale Malattie Metaboliche” Laboratory (G.Gaslini Institute) for providing us with specimens from the “Cell line and DNA bank from patients affected by Genetic diseases” Biobank - Telethon Genetic Biobank Network (project no. GTB07001A).

This study was funded by grants from Fondazione Telethon (Tiget Core Grant 2016-2021, project D2) and European Leukodystrophy Association (grant ELA 2016-010I3) to A.G.

R.D.M was supported by Fondazione Telethon (Tiget Core Grant 2016-2021, grant E5), a Career Development Award from Human Frontier Science Program (HFSP), an Advanced Research Grant from the European Hematology Association (EHA), “Pilot and Seed Grant 2015” from the San Raffaele Hospital, a Hollis Brownstein Research Grant from Leukemia Research Foundation (LRF), the Interstellar Initiative on Healthy Longevity from NYAS and AMED and by AIRC (MFAG 2019 - ID. 23321 project).

D.G was supported by the IBSA Foundation postdoctoral fellowship.

E.M was supported by a FCSR (Fondazione Centro San Raffaele) postdoctoral fellowship.

E.M., A.Ce., and L.d.V. conducted this study as partial fulfillment of their international Ph.D. in Molecular Medicine, Vita-Salute San Raffaele University.

## Author contribution

Conceptualization, A.G. and E.M.

Methodology, E.M, A.B, R.dM, S.M

Investigation, E.M., A.Ce, A.Ca., F.M., D.G., L.d.V., V.M., F.S., M.P, L.S.

Writing –Review & Editing, A.G., E.M, R.dM., A.B

Funding Acquisition, A.G.

Resources, S.M., A.B., R.dM., F.S.

Supervision, A.G., R.dM., A.B.

## Methods

Reagents generated in this study are available upon request from the Lead Contact with a completed Materials Transfer Agreement.

The lipidomic dataset generated during this study is uploaded at MetaboLights, weblink https://www.ebi.ac.uk/metabolights/MTBLS1501. The public release date is set to Feb 15 00:00:00 GMT 2021.

### Generation of iPSCs from human fibroblasts

Human dermal fibroblasts from healthy donors (ND1 and ND2) were purchased from Life technologies. Fibroblasts from GLD patients (GLD1, GLD2, GLD3, GLD4, and GLD5) and of a non-affected relative of GLD1 (ND3) were obtained from the Cell Line and DNA Bank of Patients affected by Genetic Diseases (Gaslini Institute, Genova, Italy). Human fibroblasts were cultured in Dulbecco’s modified Eagle’s medium (DMEM) supplemented with 10% fetal bovine serum (FBS), L-glutamine, and penicillin/streptomycin in an humidified atmosphere, 5% CO_2_ at 37°C. Fibroblasts at passages (p) 4-5 (GLD1, GLD2, GLD3, ND1, ND2) and p10-11 (GLD4 and GLD5, ND3) were used for reprogramming.

Reprogramming was outsourced to BioERA (Padova, Italy). Reprogramming factors were delivered as modified mRNAs (mmRNAs) taking advantage of a microfluidic system (Luni et al., 2016). The StemMACS mRNA Reprogramming Kit (Miltenyi Biotec, Bergisch Gladbach, Germany) was used for mRNA transfection according to the manufacturer instructions. Colonies positive for NANOG and TRA-1-60 were catalogued as iPSCs with progressive numbering. Six randomly selected colonies/donor were picked and expanded for 2 passages. Cells were then frozen; two frozen vials/donor were sent to SR-Tiget. The iPSCs obtained from BioERA were thawed and cultured on matrigel-coated (BD Biosciences, Erembodegen, Belgium) plastic dishes in StemMACS™ iPS-Brew XF medium (Miltenyi Biotech) in an humidified atmosphere, 5% O_2_, 5% CO_2_ at 37°C. The medium was changed daily. For routine passage, cells were detached in PBS containing 0,5 mM EDTA and split at 1:3 to 1:10 ratio. Accutase (Sigma-Aldrich, Saint Louis, USA) was used for single cell dissociation of iPSC colonies.

Human cells were used according to the guidelines on human research issued by the San Raffaele Scientific Institute’s ethics committee (protocol TIGET-HPCT).

### Karyotype analysis

Karyotyping was performed as previously described (Meneghini et al., 2017). Briefly, iPSC cultures were treated with KaryoMAX colcemid 0,1 mg/ml (Life Technologies, Carlsbad, USA) for 2 h at 37°C. After hypotonic treatment with 0,075 M KCl and fixation in methanol: acetic acid (3:1), the cell suspension was dropped onto a microscopy slide and air-dried. Chromosome counts and G-banding karyotype analyses were done on metaphases stained with Vectashield mounting medium with DAPI (Vector Laboratories, Burlingame, USA). Images were analysed with Applied Imaging Software CytoVision (CytoVision Master System). We counted >10 metaphases/coverslip. The karyotype is considered normal when >90% of the counted metaphases are normal (46, XX or 46, XY).

### Embryoid Bodies formation

iPSC colonies were gently detached using a scraper, dissociated in small cell clumps using a 2ml pipette, plated in Ultra-low attachment plates (Corning, New York, USA) and grown in suspension for 10 days in EB medium: DMEM-F12+20% Knockout Serum Replacement (KSR), non-essential aminoacids, Sodium Piruvate, L-glutamine, penicillin/streptomycin and 0,1 mM β-mercaptoethanol. The resulting spheroids were allowed to reattach on matrigel-coated 6-well plates or glass coverslips to continue differentiation for additional 10 days. Finally, they were collected for RNA extraction or fixed with 4% paraformaldehyde (PFA) for immunofluorescence analyses.

### iPSCs differentiation into NPCs

For neural induction of iPSCs we applied a dual-smad inhibition protocol (Chambers et al., 2009, 2011). Briefly, iPSC colonies were detached using Accutase, suspended as single cells and plated on Matrigel-coated dishes (75,000 cells/cm^2^) in StemMACS™ iPS-Brew XF medium supplemented with 10 μM of ROCK inhibitor Y-27632 (Sigma-Aldrich). At ≈90% confluence (usually 2 days after plating), culture medium was replaced with KSR medium (Chambers et al., 2009, 2011) supplemented with 200 ng/ml of recombinant human Noggin (R&D Systems, Minneapolis, USA) and 10 μM of SB431542 (Sigma-Aldrich). The medium was changed daily for the next 3 days. At day 4, KSR medium was gradually switched to N2 medium (Chambers et al., 2009, 2011) containing Noggin and SB431542 (75-25%, 50-50%, 25-75% every two days). After 10 days, cells were detached using Accutase and plated on Matrigel-coated dishes in N2 medium supplemented with 20 ng/ml bFGF, 20 ng/ml EGF and 10 μM Y-27632. In these conditions, the newly generated NPCs can be expanded for 4-6 passages. For all the differentiation experiments we used NPCs at passages 2-3.

### Assessment of long-term cell proliferation

iPSCs or NPCs were dissociated with Accutase and plated at a density of 30,000 or 50,000 cells/cm^2^, respectively, in the appropriate medium supplemented with 10 μM Y-27632. After 4 days, or in any case at full cell confluence, cells were treated with Accutase and counted. Cells were then replated at the initial density. The procedure was repeated for at least 4 passages.

### iPSC-NPCs differentiation into mixed neuronal/glial cultures

We applied a published protocol (Frati et al., 2018) with some modifications. Cells were detached using Accutase and plated on Matrigel-coated dishes (20,000 cells/cm^2^) in N2 medium with 20 ng/ml bFGF, 20 ng/ml EGF and 10 μM Y-27632 (d0). Two days after plating and for the subsequent 4 days we gradually replaced N2 medium with increasing amount of Glial Differentiation Medium (GDM) containing 10 ng/ml PDGF-AA, 10 ng/ml NT3, 10 ng/ml IGF-1, 5 ng/ml HGF, and 60 ng/ml T3. From d4 to the end of the differentiation protocol (d24) medium was changed every other day. From d14 to day 24 cells were kept in Glial Maturation Medium (GMM) in absence of growth factors and supplemented with 20 μg/ml of ascorbic acid (Sigma-Aldrich).

### Lentiviral vector-mediated *hGALC* gene transfer in iPSCs

We used a previously described (Meneghini et al., 2016) third-generation lentiviral vector (LV) encoding the human (h)GALC cDNA tagged with the myc peptide, under control of the human phosphoglycerate kinase (PGK) promoter. The titer of the LV batch was 1.6×10^9^ TU/ml. For cell transduction, iPSC colonies were dissociated at single cells using Accutase and plated on Matrigel-coated 6 multi-well plates (10,000 cells/cm^2^) in StemMACS™ iPS-Brew XF medium supplemented with 10 μM Y-27632. After 6 hours, cells were incubated with LV.hGALC vector at different MOI (0-100) in StemMACS™ iPS-Brew XF medium additioned with 10 μM Y-27632 and 8 μg/ml Polybrene. After an obvernight incubation, we added fresh culture medium. Cells were passaged as they reached confluency, as described. Transduced cells were used for the experiments after 3-4 subculturing passages, in order to dilute the non-integrated LV genome.

### Quantification of Psychosine

Psychosine quantification was performed on frozen cell pellets by TSQ Quantum AM mass spectrometer (Thermo Fisher Scientific, Waltham, USA) as previously described (Lattanzi et al., 2014).

### GALC activity assay

Specific beta-galactocerebrosidase activity was measured by hydrolysis of artificial fluorogenic substrate 4-methylumbelliferone-beta-galactopyranoside in presence of the competitive inhibitor of beta-galactosidase AgNO_3_, as previously described (Martino et al., 2009).

### PLA2 activity assay

Quantification of Phospholipase A2 (PLA2) activity was performed using the EnzChek Phospholipase A2 Assay Kit (Thermo Fisher). 50 μl of cell lysate/well, corresponding to 20 μg total proteins, was mixed with 50 μl of substrate cocktail to initiate the catalytic reaction. After 10 minutes, fluorescence was measured using a Victor3 Multilabel Counter (Perkin Elmer, Waltham, USA) at 360 nm (excitation) and 460 nm (emission). The specific enzymatic activity was calculated for each sample and normalised for protein content. Values are expressed as relative fluorescent units (RFU)/μg of protein/min.

### SA-β-Galactosidase assay (X-gal staining)

Induction of senescence during NPCs differentiation was assessed with a senescence β-galactosidase (SA-β-Gal) Staining Kit (Cell Signalling Technology, Danvers, USA) according to manufacturer’s instructions. In detail, NPCs were plated on coverslips and fixed at day 0, day 7 and day 14 days of differentiation with4% PFA for 10 minutes at RT and then incubated for 24 hours with the SA-β-gal staining solution (pH=6.0) to reveal SA-β-gal activity. Nuclei were then stained with Hoechst. Images were acquired using a Nikon Eclipse Ni microscope, visualized and analysed using the ImageJ software. Cells displaying blue signal in the cytosol were counted as positive. At least 850 nuclei/coverslip/timepoint were used for quantification.

### Proliferation analysis (EdU staining)

EdU (5-ethynyl-2′-deoxyuridine), supplied with Click-iT EdU Alexa Fluor 647 Imaging Kit (Thermo-Fisher Scientific), was diluted in DMSO to a final concentration of 10 mM and kept at −20°C. Cells were incubated with 2μM EdU overnight in culture. 2×10^5^ cells per sample were transferred to cytometry tubes and then washed with PBS 1% BSA and fixed with 100 μL of Click-iT fixative for 15 minutes. Cells were washed again with PBS 1% BSA and permeabilized with 100 μL of 1X Click-iT saponin-based solution for 15 minutes. Detection of EdU-DNA was performed by incubating cells with 500 μL of Click-iT Plus reaction cocktail for 30 minutes at RT protected from light. Cells were subsequently washed with PBS 1% BSA and fluorescence was measured by flow cytometry.

### High Throughput live cell Microscopy

Determination of lysosomal area was performed using a high content imaging platform in the Advanced Light and Electron Microscopy Bio-Imaging Centre of San Raffaele Scientific Institute (ALEMBIC) using the ArrayScan XTI HCA Reader from Thermo Fisher Scientific. Cells were plated on matrigel-coated 96 Greiner Sensoplate glass bottom multiwell plates (Sigma Aldrich) (30,000 cells/well). The day after plating, cells are incubated with 50 nM LysoTracker Red DND-99 (Thermo Fisher) diluted in culture medium, for 30 minutes at 37°C. Nuclear staining, necessary for the instrument acquisition and analysis, was performed using Hoechst (5 min at room temperature, 10 μg/ml final concentration). Cells are then washed with Krebs-Ringer-Hepes-bicarbonate buffer (KRH) and maintained in the same buffer for the entire duration of the instrument acquisition. Thirty fields were acquired for each well; images were analysed with the Thermo Scientific HCS Studio Cell Analysis Software

### Immunofluorescence

Cells are seeded in 48-multiwell plates on Matrigel-coated 10 mm round sterile glass coverslips, fixed in 4% PFA, rinsed with PBS, incubated with blocking solution containing 10% normal goat serum (NGS) + 0.1% Triton X-100 in PBS for 45 minutes and incubated overnight at 4°C with primary antibodies diluted in blocking solution. After 3 washes (10 minutes each), cells are incubated with species-specific fluorophore-conjugated secondary antibodies diluted in 1% NGS in PBS for 1 hour at room temperature. Cell nuclei are counterstained with Hoechst (10μg/ml, diluted in PBS). Finally, coverslips are mounted on glass slides using Fluorsave mounting medium (Merck Millipore). A secondary-only control was included in every experiment to confirm signal specificity. Images were obtained with a Nikon Eclipse Ni microscope, visualized and analysed using the ImageJ software.

To evaluate DDR activation during NPCs differentiation, cells were fixed with 4% PFA for 20 min at RT and then permeabilized with 0.3% Triton-X100 in PBS for 10 min. After washing, coverslips were blocked in PBG (0.2% cold water fish gelatine, 0.5% BSA in PBS) for 30 min. Cells were then stained with primary antibodies 1 hour at RT followed by secondary antibodies. After DAPI staining (Sigma-Aldrich), slides were mounted with Aqua-Poly/mount (Polysciences Inc, Warrington, USA). Images were obtained with Leica TCS SP5 confocal laser microscope and visualized with Leica Application Suite (LAS) software (Leica Microsystems, Wetzlar, Germany). Quantification of DDR activation in immunofluorescence images was conducted by counting colocalizing foci for 53BP1 and γH2A.X.

Primary and secondary antibodies used:

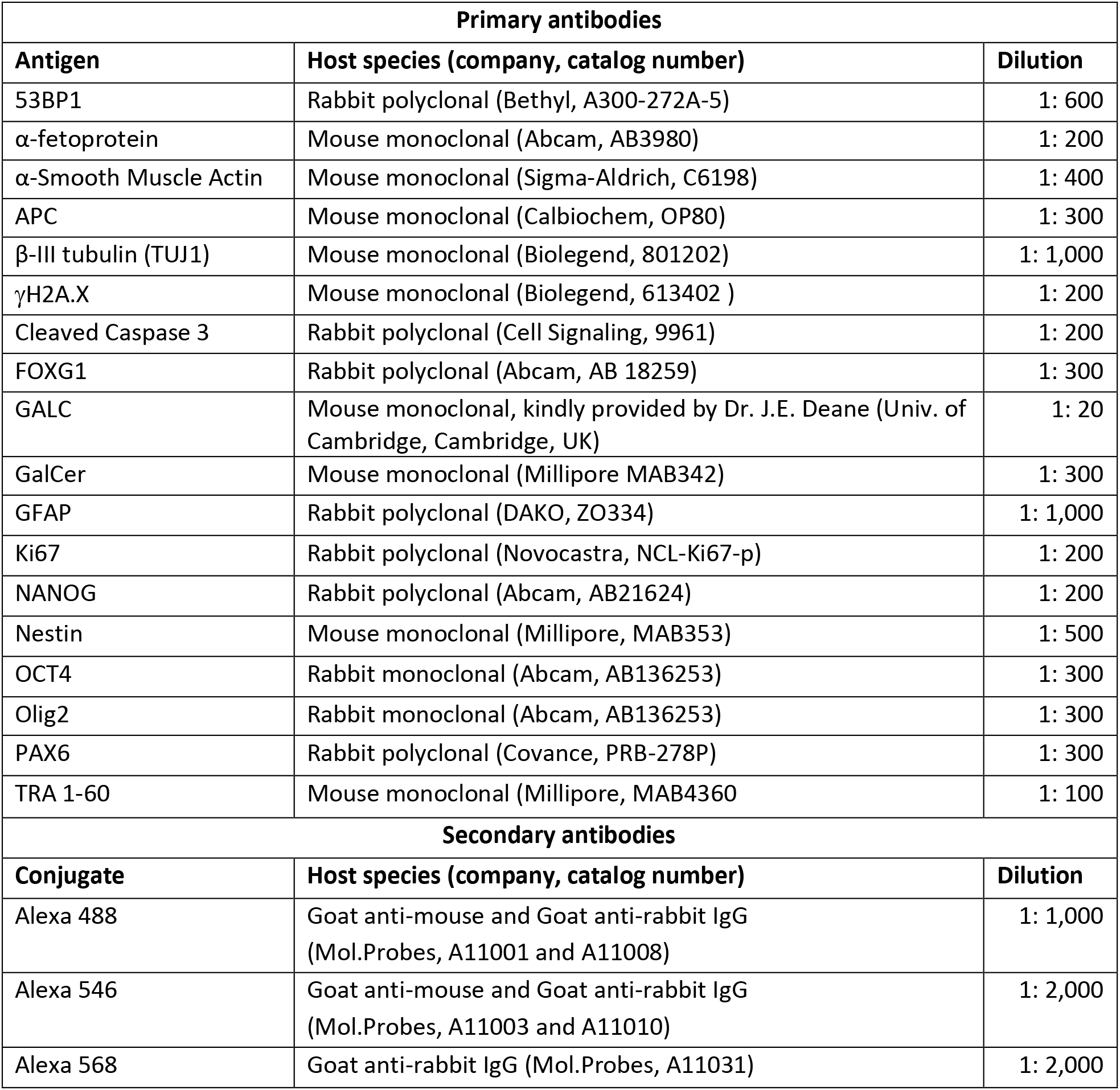

### Western blot analysis

Cell pellets were lysed using RIPA buffer (50 mM Tris–HCl pH 7,4, 150 mM NaCl, 0,5% sodium deoxycholate, 0,1% SDS, 2 mM EDTA) containing protease and phosphatase inhibitors (Roche, Basel, Switzerland). Protein content was measured using the Bradford reagent (Bio-Rad, Hercules, USA) with bovine serum albumin (BSA) as the reference standard. SDS-PAGE of protein extracts was performed using Novex NuPAGE SDS-PAGE system (Thermo Fisher Scientific) according to manufacturer instructions. Proteins were transferred on PVDF membranes (Millipore, Burlington USA) for 1 hour at 400mA.

Primary and secondary antibodies used:

| Primary antibodies |  |  |
| --- | --- | --- |
| Antigen | Host species (company, catalog number) | Dilution |
| α-tubulin | Mouse monoclonal (Sigma-Aldrich, T6074) | 1: 5,000 |
| Acetylated α-tubulin | Mouse monoclonal (Sigma-Aldrich, T7451) | 1: 20,000 |
| β-Actin | Rabbit polyclonal (Santa Cruz, sc-1616r) | 1: 10,000 |
| β-Actin | Mouse monoclonal (Sigma-Aldrich, A2228 ) | 1: 10,000 |
| β-III tubulin (TUJ1) | Mouse monoclonal (Biolegend, 801202) | 1: 6,000 |
| Calnexin | Rabbit polyclonal (Sigma-Aldrich, c4731) | 1: 10,000 |
| GALC | Rabbit polyclonal (Abcam, AB137750) | 1:3,000 |
| GFAP | Mouse monoclonal (Millipore, MAB3402) | 1: 100,000 |
| LAMP1 | Rabbit polyclonal (Abcam, AB24170) | 1: 6,000 |
| MYC | Rabbit polyclonal (Abcam, AB9106) | 1: 15,000 |
| Neurofilament-M (RMO 14.9) | Mouse monoclonal (Cell Signaling, 2838) | 1: 1,000 |
| OLIG1 | Rabbit polyclonal (Millipore, AB5591) | 1: 800 |
| p21 | Rabbit monoclonal (Cell Signaling, 2947) | 1: 1,000 |
| Polyglutamylation Modification | Mouse monoclonal (Adipogen, AG-20B-0020) | 1: 5,000 |
| SMI 311 | Mouse monoclonal (Sternberg, SMI-311) | 1:1000 |
| SMI 312 | Mouse monoclonal (Covance, SMI312R) | 1:1000 |
| Secondary antibodies |  |  |
| Conjugate | Host species (company, catalog number) | Dilution |
| HRP | Goat anti-mouse IgG (Chemicon, AP124P) | 1: 5,000 |
| HRP | Goat anti-rabbit IgG (Millipore, 12-348) | 1: 10,000 |

### Molecular analysis

#### Sequencing of patient’s mutations

Genomic DNA was extracted from iPSCs pellets using Maxwell cell DNA purification Kit (Promega), following the manufacturers’ instructions. DNA concentration was determined by 260/280nm optical density (OD) reading on a NanoDrop ND-1000 Spectrophotometer (NanoDrop, Pero, Italy). To confirm the presence of the expected mutations in the *GALC* gene of GLD patients-derived cells, the regions of interests were amplified by PCR, purified from agarose gels using the Wizard SV Gel and PCR Clean-Up System (Promega, Madison, USA) and sent for sequencing (to GATC Services-Eurofin Genomics, Ebersberg, Germany). Sequences were checked using the Snapgene software. Three primer PCR to confirm 30kb deletion in GLD1 and GLD3 cells was performed as described by (Liu et al., 2011). Primers used for PCR amplification and sequencing are the following:

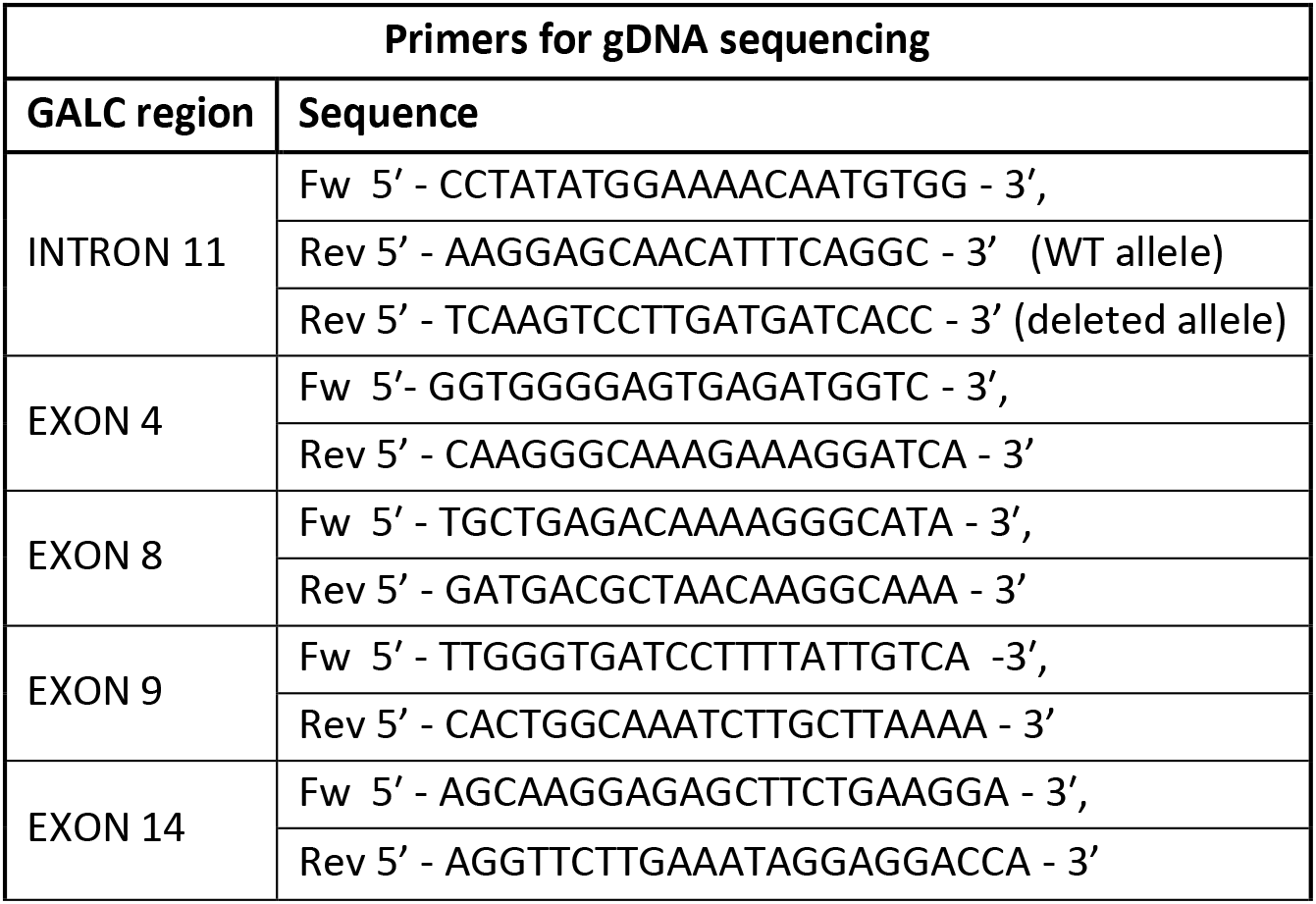

#### Gene expression analysis: RT-PCR; SYBR green and TaqMan RT-PCR

Total RNA was extracted using the RNeasy Mini Kit (Qiagen). RNA was quantified by 260/280nm optical density (OD) reading on a NanoDrop ND-1000 Spectrophotometer (NanoDrop, Pero, Italy). mRNA reverse transcription (RT) is performed with QuantiTect reverse transcription kit (Qiagen, Hilden, Germany).

<u>RT-PCR analyses</u> were performed by using FastStart Taq DNA Polymerase, PCR KIT (Roche). For each reaction, 100 ng of template cDNA and 1 mM of primers were used. PCR products were resolved by electrophoresis in a 1,5% agarose gel.

<u>Quantitative real time PCR reactions</u> were performed in Optical 96-well Fast Thermal Cycling Plates (Primo 96-well plates, Euro Clone, Pero, Italy) on Viia7 quantitative real time PCR system (Applied Biosystem, Foster City, USA), using the following thermal cycling conditions:

– SYBR green qPCR: one cycle at 95°C for 2 minutes, 40 cycles at 95°C for 10 seconds and 60°C for 30 seconds; each sample is run in duplicate in a total volume of 12.5 μl/reaction, containing 6.25 μl 2X Fast SYBR Green Master Mix (Qiagen), 10 ng of template cDNA, and10μM of custom primers were used. As internal reference for normalization GAPDH was used
– TaqMan qPCR: one cycle at 95°C for 5 minutes, 40 cycles at 95°C for 50 seconds and 60°C for 1 minute. Each sample is run in duplicate in a total volume of 12.5 μl/reaction, containing 6.25 μl 2X EagleTaq Universal Master Mix (Roche), 1μl of sample cDNA and 0.63μl of Probe+Primers mix (TaqMan Gene Expression Assays, Thermo Fisher). The Viia7 software was used to extract raw data. Relative expression of mRNA for the target genes was calculated using the comparative Ct (ΔΔCt) method, with a housekeeping gene as control. Raw data (Ct values) were analyzed according to the comparative Ct method. For gene expression data, analyses were performed on 2(-^ΔCt^) values relative to housekeeping gene or plotted as 2^−ΔΔCt^ (fold change), as specified in figure legends. Primers and probes used for gene expression analyses are the following:

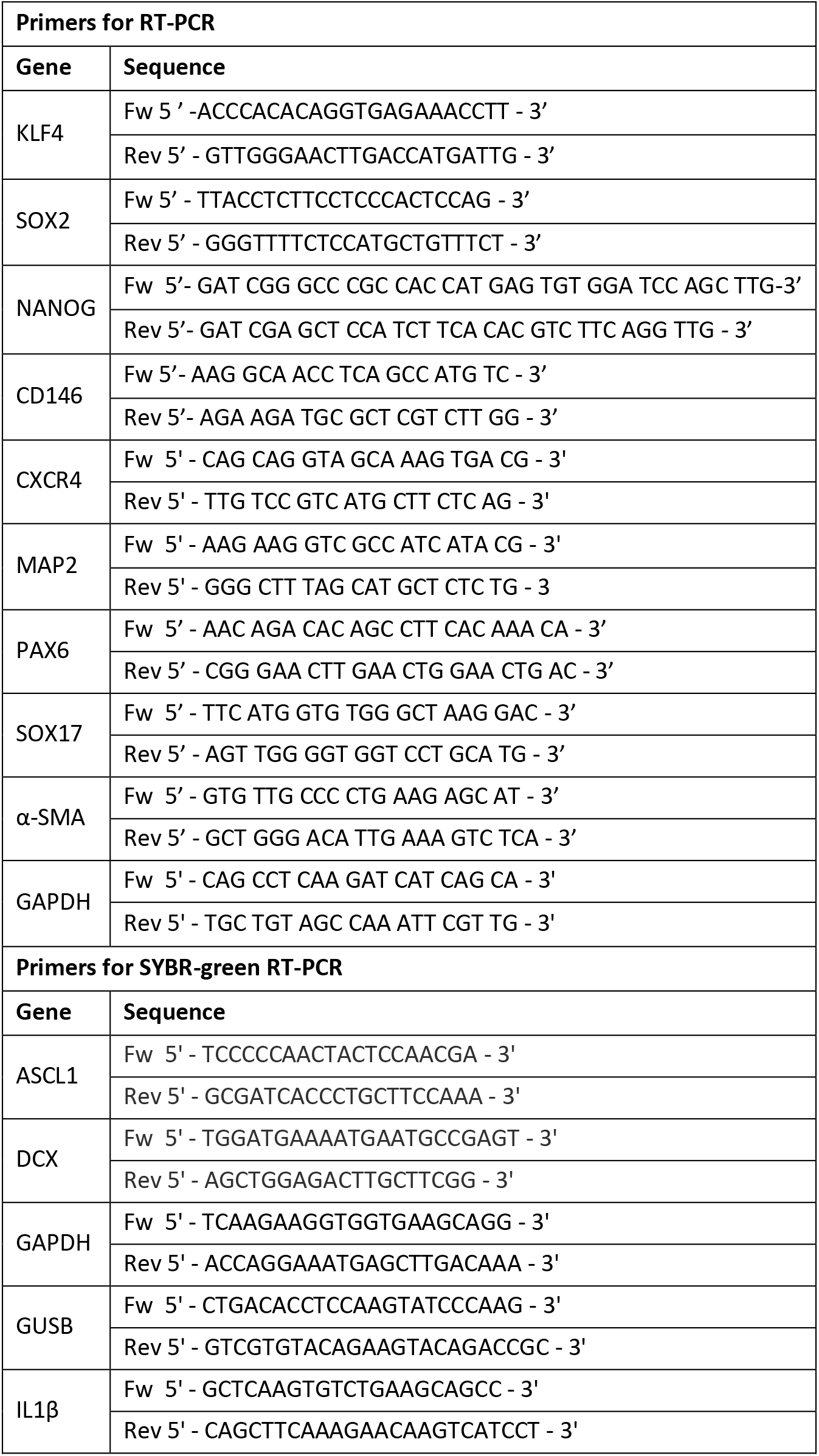
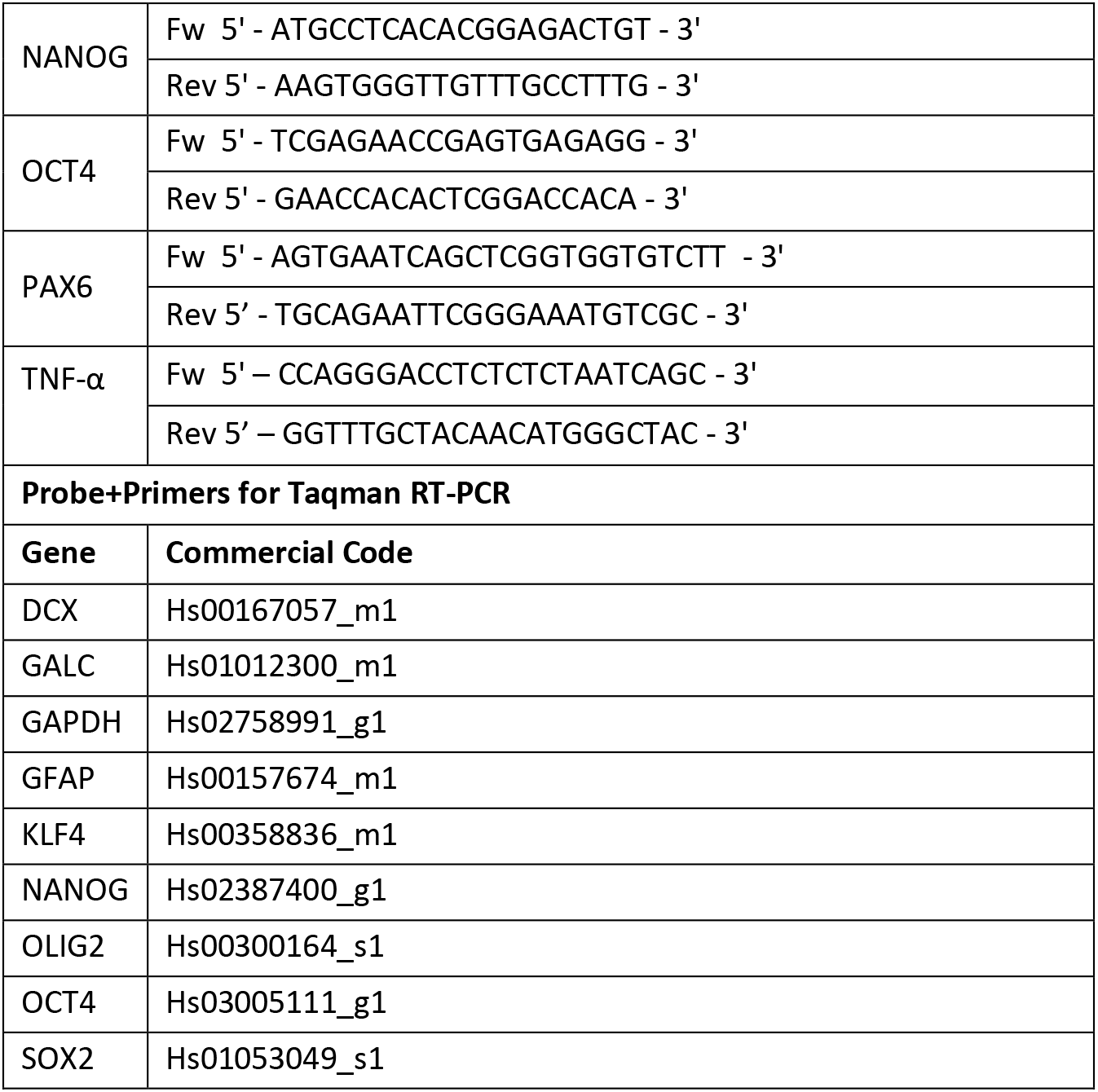

### Determination of vector copy number (VCN)

Genomic DNA was extracted from cell pelletes using Maxwell cell DNA purification Kit (Promega) or QIAamp DNA Micro Kit (Quiagen), depending on the number of cells. The number of integrated vector copies/genome was quantified by TaqMan analysis, as previously described (Lattanzi et al., 2010). Briefly, quantitative PCR was performed on 50 ng of template DNA using probes and primers that anneal to the PSI (ψ) sequence of the LV backbone and to a fragment of the human telomerase gene. A standard curve was set up with gDNA extracted from a human lymphoblast cell line (CEM) carrying 1 copy of integrated LV.GFP, validated by Southern blot analysis. The standard curve, based on different dilutions of CEM gDNA (from 0,32 ng to 40 ng) was used for both LV and telomerase amplification. Reactions were carried out in a total volume of 25 μl, on Viia7 quantitative real time PCR system (Applied Biosystem). Eventually, VCN is calculated as: (ng LV/ng endogenous DNA) × (number of LV integrations in the standard curve) × cell ploidy. Primers and probes used for VCN analyses are the following:

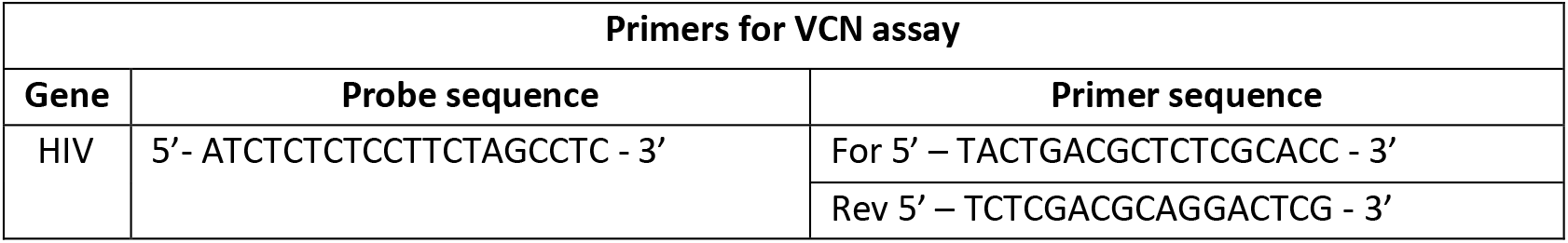
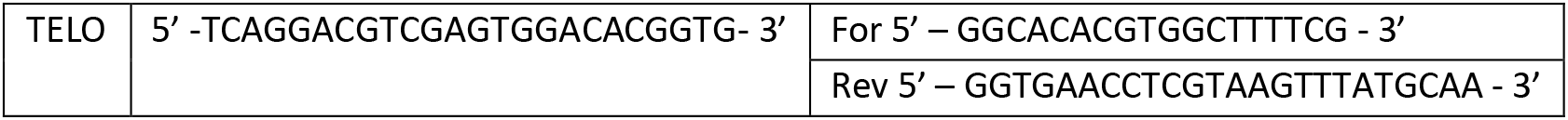

### Lipidomic analyses

#### Lipid extraction

Cell pellets were resuspended in 150 mM Ammonium bicarbonate and passed through a 26G syringe needle to prepare cell lysate. Lipids were extracted starting from an equivalent of 10 μg of proteins, using a 2 steps extraction protocol (Folch method) with methanol and chloroform in different proportions (FOLCH et al., 1957). Organic phase fractions were then dried out and resuspended in 50 μL of CH3OH for subsequent analysis. Before extraction, samples were spiked in with 16 internal standards: PC (12:0/13:0) 40 pmol, PE (12:0/13:0) 52 pmol, PG (12:0/13:0) 7.5 pmol, PS (12:0/13:0) 43 pmol, PI (12:0/13:0) 54 pmol, Cer (d18:1/25:0) 100 pmol, CE(19:0) 100 pmol, GlcCer (d18:1/12:0) 50 pmol, LacCer (d18:1/12:0) 50 pmol, Sphinganine (d17:0) 50 pmol, Sphingosine-1-P (d17:1) 100 pmol, Sphingosine (d17:1) 50 pmol, Galactosyl(ß) Sphingosine-d5 20 pmol, d5-TG ISTD Mix I 20 pmol, d5-DG ISTD Mix I 20 pmol, cholesterol (d7) 800 pmol.

#### Protein quantification

Proteins were extracted form 20 μL of ammonium bicarbonate resuspended fractions by adding 5 μL of lysis buffer (10% NP40, 2%SDS in PBS) and quantified by BCA protein assay kit (ThermoFisher Scientific, 23225).

#### Lipid profiling data acquisition

Lipids extracts were diluted 1:5 with 95% mobile phase A (CH3CN:H2O 40:60; 5mM NH4COOCH3; 0.1% FA) plus 5% mobile phase B (IPA:H2O 90:10; 5mM NH4COOCH3; 0.1% FA) and 1 μL injected on a liquid chromatography system nLC Ekspert nanoLC400 (Eksigent, 5033460C; Singapore) coupled with a Triple TOF 6600 (AB Sciex; Singapore). Chromatography was performed using an in house packed nanocolumn Kinetex EVO C18, 1.7 μm, 100 A, 0.75 × 100 mm. The gradient elution at flow rate of 150 nL/min was initially started from 5 % of mobile phase B, linearly increased to 100 % B in 5 min, maintained for 45 min, then returned to the initial ratio in 2 min and maintained for 8 min. Positive mode acquisition in MS was performed with the following parameters: mass over charge (m/z) range 100– 1700, T source 80C, Ion Spray Voltage 2000, Declustering Potential (DP) 80, fixed collision energy 40 V (+). For IDA analysis (Top 8), range of m/z was set as 200–1800 in positive ion mode, target ions were excluded for 20 sec after 2 occurrences (Analyst TF 1.7.1).

#### Data processing and analysis

Lipidview workstation (version 1.3 beta, AB SCIEX, USA) was used for lipids identification and quantitation. Lipid identification was based on exact mass, retention time, and MS/MS pattern. Lipid species based on precursor fragment ion pairs were determined using a comprehensive target list in LipidView (Sciex). Lipid species identification was performed using; the mass tolerance of 0.05 in MS and 0.02 in MS/MS, s/n of 3 and % peak intensity > 0 for positive ion mode.

Lipid classes included for statistics and downstream analysis were: cholesterol ester (CE), sphingomyelin (SM), diacylglycerol (DAG), triacylglycerol (TAG), ceramide (Cer) phosphatidylcholine (PC), phosphatidylethanolamine (PE), phosphatidylglycerol (PG), phosphatidylinositol (PI), phosphatidylserine (PS) and lysophosphatidylcholine (LPC), lysophosphatidylethanolamine (LPE), lysophosphatidylglycerol (LPG), lysophosphatidylinositol (LPI), lysophosphatidylserine (LPS), hexosylceramide (HexCer), dihexosylceramide (Hex2Cer), trihexosylceramide (Hex3Cer), sulphatides (SGalCer), ceramide-phosphate (CerP), in positive mode.

### Statistics

Data were analysed with Graph Pad Prism version 8 for Macintosh and expressed as the mean ± standard error of the mean (SEM). The number of samples and the statistical tests and post-tests used are specified in the legends to each figure. p<0.05 was considered statistically significant.

For lipid analyses, statistical analysis was performed using Metaboanalyst 4.0 web tool https://www.metaboanalyst.ca/MetaboAnalyst/faces/home.xhtml (Chong et al., 2018).

