## Supplementary material for "Human iPSC-based neurodevelopmental models of globoid cell leukodystrophy uncover patient- and cell type-specific disease phenotypes"

#### Supplementary Tables and Figures

**Table S1. ND and GLD fibroblast cell lines used in this study**

The table summarizes the origin of normal donor (ND) and patient-specific (GLD) fibroblast cell lines, and the disease-causing mutation (including the disease variant, the allele's reference and the mutated protein). *GALC* mutations are reported according to current nomenclature guidelines, ascribing the A of the first ATG translational initiation codon as nucleotide +1 (http://www.hgvs.org/mutnomen). We provide bibliographic reference (when available) of studies reporting the specific mutations and the predicted features of the mutated protein.

| ID | Provider (code) | Sex | Mutation | Protein | Disease form | Reference |
|---|---|---|---|---|---|---|
| **GLD1** | Gaslini Biobank (FFF0461990) | F | c.1161+6532_polyA + 9Kbdel | Truncated protein | Infantile | Selleri et al, J Neurol. 2000 Wenger et al., 1997 Tappino et al., 2010 |
| **GLD2** | Gaslini Biobank (FFF0112003) | F | c.379C>T (nonsense mutation in EXON 4) | p.R127X mRNA instability | Infantile | Tappino et al., 2010 |
| | | | c.1657G>A (missense mutation in EXON 14) | p.G553R Severe misfolding | | De Gasperi et al, 1999 |
| **GLD3** | Gaslini Biobank (FFF0062002) | M | c.941A>G (missense mutation in EXON 9) | p.Y314C | Infantile | De Gasperi et al, 1996 Shin et al., 2016 |
| | | | c.1161+6532_polyA + 9Kbdel | Truncated protein | | |
| **GLD4** | Gaslini Biobank (FFF0332007) | M | c.388G>A (missense mutation in EXON 4) | p.E130K Severe misfolding | Infantile | Lissens et al 2007 Tappino et al 2010 Berardi et al 2014 |
| | | | c.884A>C (missense mutation in EXON 8) | p.N295T Loss of stabilizing hydrogen bonds | | Berardi et al 2014 Wenger et al 1997 Spratley et al 2016 |
| **GLD5** | Gaslini Biobank (FFF0042010) | F | c.1657G>A (missense mutation in EXON 14) | p.G553R Severe misfolding | Infantile | De Gasperi et al 1999 Berardi et al. 2014 Tappino et al., 2010 Deane et al. 2011 |
| **ND1** | Life Technologies HDF adult (C0135C) | F | na | wt | na | na |
| **ND2** | Life Technologies HDF neonatal (C0045C) | M | na | wt | na | na |
| **ND3** | Gaslini Biobank (FFF0171991) | M | c.1161+6532_polyA + 9Kbdel (heterozygote) | wt/truncated protein | na | na |

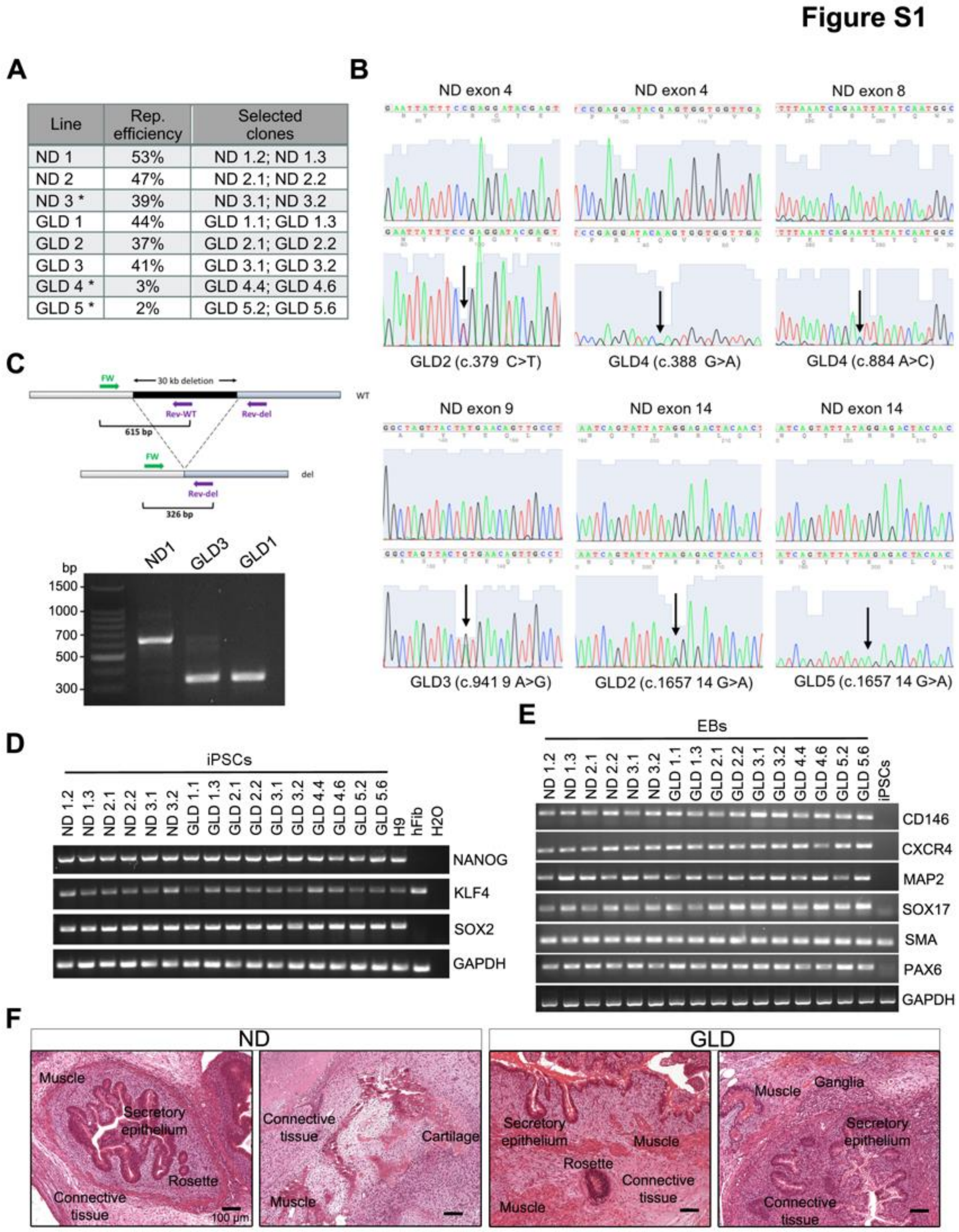

**Figure S1. Generation and characterization of ND and GLD iPSCs**

A) List of ND and GLD iPSC clones selected for characterization of pluripotency. Reprogramming efficiency was calculated as the percentage of primary iPSC colonies generated on the number of plated fibroblasts. Asterisks: fibroblasts at high passages (p10-11) at the time of reprogramming – see also Methods.

B) Sanger sequencing performed of GLD iPSCs DNA highlights the mutations in exon 4 (GLD2 and GLD4), exon 8 (GLD4), exon 9 (GLD3), and exon 14 (GLD2, GLD5) of the human *GALC* gene. The sequences retrieved by ND1.2 iPSCs DNA are shown for comparison.

C) Schematic representation of three primers PCR approach performed to verify the presence of the 30kb deletion in the h*GALC* gene of patient's derived iPSCs. The amplification product demonstrates the presence of the 30 kb deletion in iPSCs from patient GLD3 (heterozygote) and patient GLD1 (homozygote).

D) RT-PCR analysis showing mRNA expression of NANOG, KLF4 and SOX2 in the selected ND and GLD iPSC clones. Embryonic stem cells (line H9; kindly provided by Dr. A. Ditadi) were used as positive control; human fibroblasts (ND1) were used as negative control. Endogenous KLF4 is expressed in all cell types. GAPDH was used as housekeeping gene.

E) RT-PCR analysis showing mRNA expression of endodermal (SOX17, CXCR4), mesodermal (CD146, $\alpha$-SMA), and ectodermal markers (PAX6, MAP2) in iPSC-derived EBs and in one iPSC line (ND1, last lane). GAPDH was used as housekeeping gene.

F) Representative images of ND and GLD iPSC-derived teratoma sections stained with haematoxylin/eosin show the presence of endodermal, mesodermal and endodermal tissues. Scalebar= 100 $\mu$m.

#### Figure S2

**A**

| SAMPLE | NUMBER OF METAPHASES ANALYSED | KARYOTYPE | DESCRIPTION |
|---|---|---|---|
| ND 1.1 p33 | 10 | 47,XX+14 | Trisomy of chromosome 14 (100% of metaphases) |
| **ND 1.2 p15** | **10** | **46,XX** | **Normal (100% of metaphases)** |
| ND 1.2 p31 | 12 | 46,XX | Isochromosome 14 (>40% of metaphases) |
| **ND 1.3 p13** | **12** | **46,XX** | **Normal (100% of metaphases)** |
| ND 1.6 p15 | 10 | 46,XX | Trisomy of chromosome 14 (40% of metaphases) |
| **ND 2.1 p15** | **11** | **46,XY** | **Normal (100% of metaphases)** |
| **ND 2.2 p12** | **10** | **46,XY** | **Normal (100% of metaphases)** |
| **GLD 1.1 p24** | **10** | **46,XX** | **Normal (100% of metaphases)** |
| **GLD 1.3 p13** | **16** | **46,XX** | **Normal (94% of metaphases)** |
| GLD 1.6 p12 | 13 | 46,XX ,derX,der2 | Derivative X and 2 crhomosomes (100% of metaphases) |
| **GLD 5.2 p16** | **10** | **46,XX** | **Normal (100% of metaphases)** |
| GLD 5.2 p39 | 21 | 46,XX | Trisomy of chromosomes 14 and 20 (>20% of the metaphases) |
| **GLD 5.6 p13** | **10** | **46,XX** | **Normal (100% of metaphases)** |

**B**

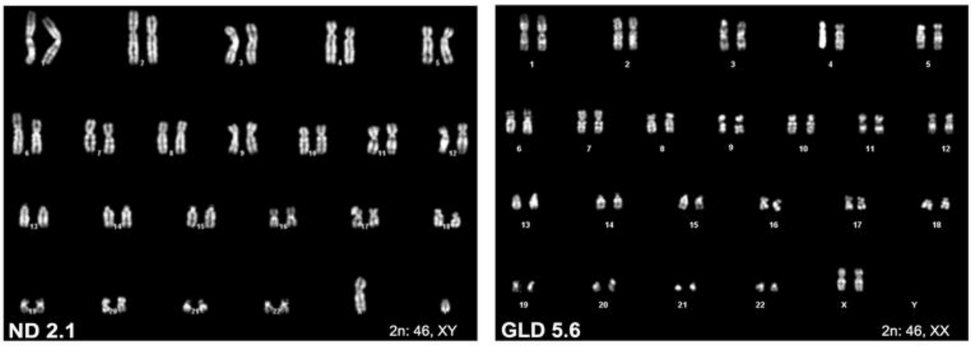

**Figure S2. Karyotype analyses of ND and GLD iPSC clones**

A) Karyotype assessed by G-banding in ND and GLD iPSC clones at different passages (p) in culture (p12-p39). Analyses were performed on at least 10 metaphases/clone. Clones with normal karyotype (highlighted in bold) were selected for further characterization.

B) Representative picture showing the normal karyotype of ND2.1 (46, XY) and GLD5.6 clones (46, XX).

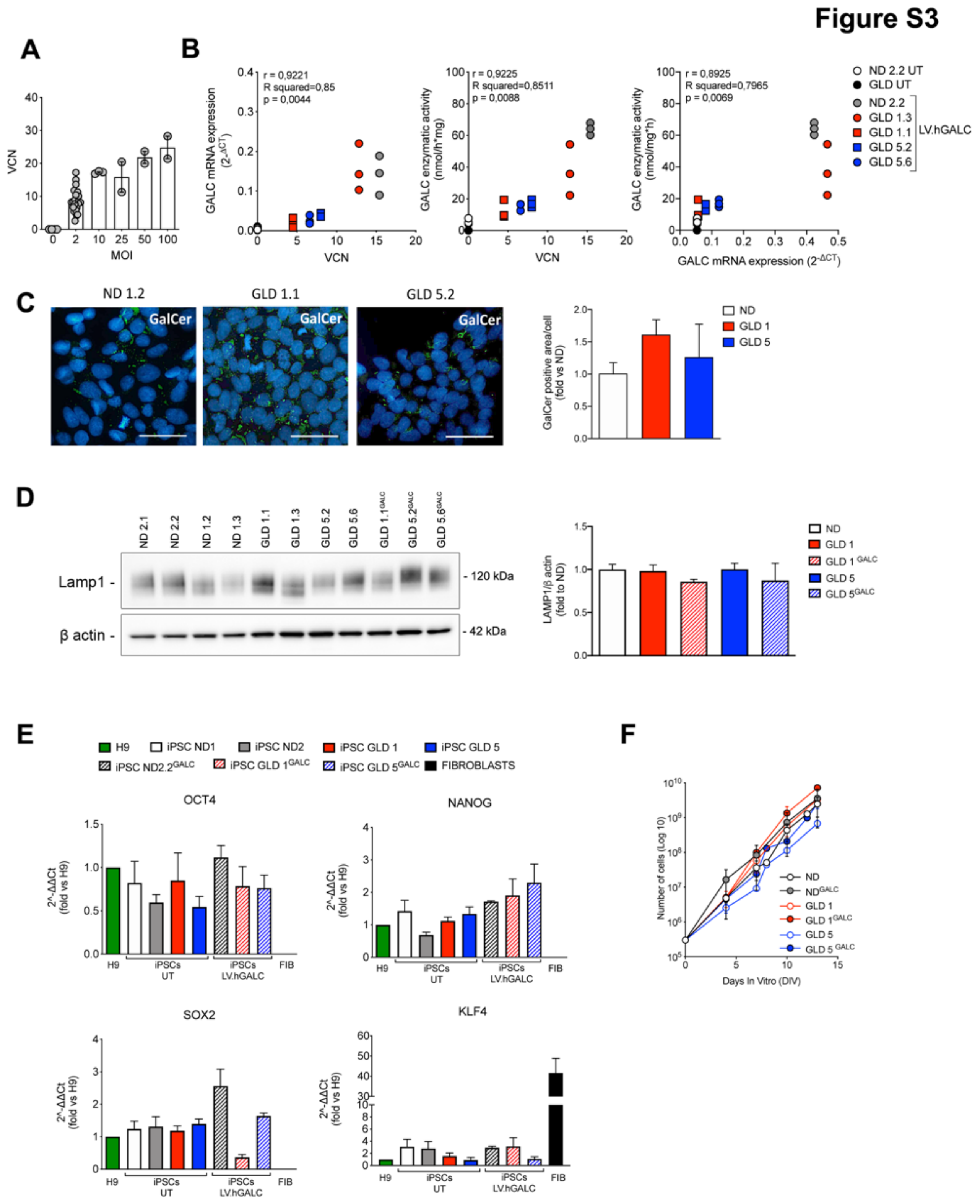

**Figure S3. Functional properties of untransduced and LV.hGALC-transduced ND and GLD iPSCs**

A) Vector Copy Number (VCN) assessed in LV.hGALC-transduced iPSCs. Multiplicity Of Infection (MOI) ranges from 0 (untransduced cells) to 100. All ND and GLD iPSC clones transduced at the same MOI have been pooled together. Clones used: ND2.2, GLD1.1, GLD1.3, GLD5.2, GLD5.6**.** n=1-4 clone/group, 2–6 replicates/clone. Data are represented as the mean ± SEM.

B) Positive correlation (Pearson r; p<0.05) between i) VCN and GALC mRNA expression; ii) VCN and GALC enzymatic activity; iii) GALC mRNA expression and GALC enzymatic activity in LV.hGALC-transduced ND and GLD iPSC clones. Untransduced ND and GLD clones are shown for comparison.

C) Representative immunofluorescence pictures of ND and GLD iPSCs expressing GalCer (green). Nuclei are counterstained with Hoechst. Scale bars=50 μm. The graph shows the quantification of the GalCer+ area normalized on the nuclear area (in pixels) and expressed as fold to ND (mean of all clones). Data are the mean+SEM; n=2 independent experiment, 3-6 replicates, 5-10 fields analyzed/sample. Clones used: ND1.2, ND1.3, ND2.1, ND2.2, GLD1.1, GLD1.3, GLD5.2, GLD5.6). Data were analyzed by one-way ANOVA followed by Dunnett's multiple comparison post-test. No significant differences between groups (p>0.05).

D) Representative western blot images and densitometry quantification showing the expression of LAMP1 protein in ND, GLD, and $GLD^{GALC}$ iPSC clones. β -actin was used as loading control. Data are represented as the mean+SEM. n= 3 replicates/clone; 1-4 clones/group. Data were analyzed by two-way ANOVA followed by Dunnett's multiple comparison post-test (control group: ND clones). No significant differences between groups (p>0.05).

E) mRNA expression (assessed by qRT-PCR) of pluripotency markers (OCT4, NANOG, SOX2, KLF4) in untransduced (UT) and LV.hGALC-transduced iPSCs. CT values for the gene of interest are normalized on GAPDH and calculated as fold change ($2^{-\Delta\Delta CT}$) on the expression of the hESC line H9. Human fibroblasts (FIB; ND1) were included as control. Fold change values of OCT4, NANOG, and SOX2 in FIB are <0.0005. Data are expressed as the mean+SEM; n= 2 independent experiments, 1-3 clones/group. Clones used: ND1.2, ND1.3, ND2.1, ND2.2, GLD1.1, GLD1.3, GLD5.2, GLD5.6, $GLD1.1^{GALC}$, $GLD5.2^{GALC}$, $GLD5.6^{GALC}$.

F) Stable expansion rate of untransduced and LV.hGALC-transduced ND and GLD iPSC lines (4-5 subculturing passages are shown in the graph; up to 13 days in vitro, DIV). Each point of the curves represents the mean ± SEM; n=2–5 independent experiments, 1-4 clones/group. Clones used: ND2.1, ND2.2, GLD1.1, GLD1.3, GLD5.2, GLD5.6, $ND2.2^{GALC}$, $GLD1.1^{GALC}$, $GLD5.2^{GALC}$, $GLD5.6^{GALC}$.

**Figure S4**

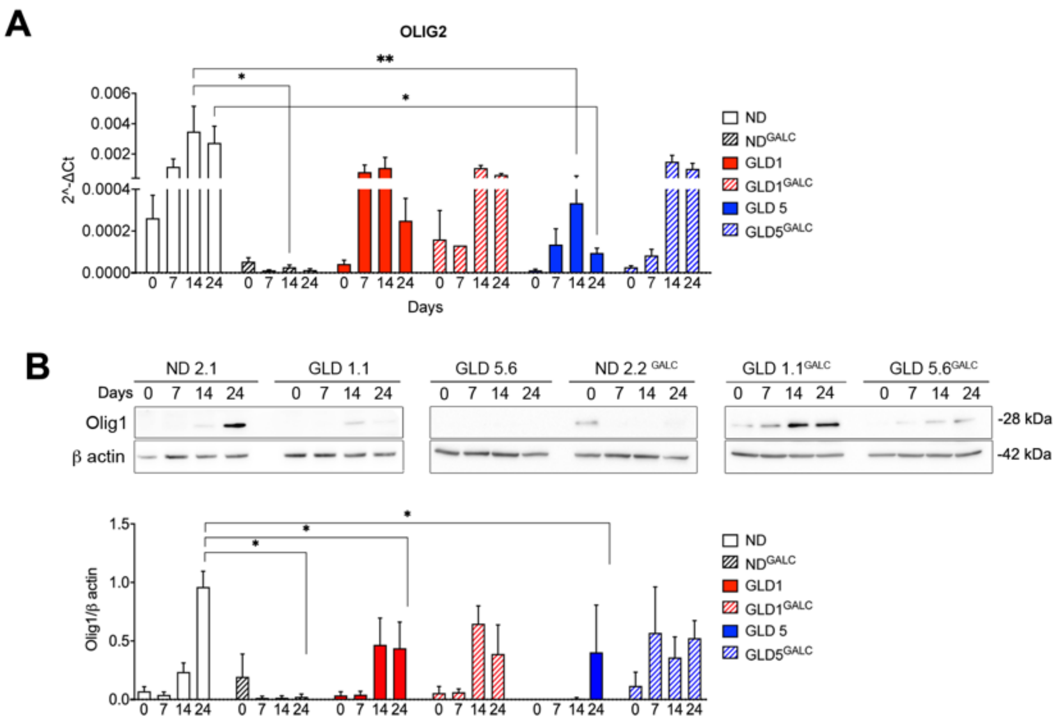

**Figure S4. Expression of oligodendroglial markers during NPC differentiation**

A) OLIG2 mRNA expression (assessed by qRT-PCR) in untransduced (plain bars) and LV.hGALC-transduced (striped bars) ND and GLD NPCs (d0) and differentiated neural progeny (d7, d14 and d24). Values are normalized on GAPDH and expressed as 2^-ΔCT (mean+SEM). Clones used: ND1.2, ND2.1, ND2.2, GLD1.1, GLD1.3, GLD5.2, GLD5.6**,** ND2.2$^{GALC}$, GLD1.1$^{GALC}$, GLD5.2$^{GALC}$, GLD5.6$^{GALC}$. n= 2-4 independent experiments; 1-3 clones/group. Data were analyzed by two-way ANOVA followed by Dunnett's multiple comparison post-test (control group: ND at the corresponding time point) $^*p < 0.05$, $^{**}p < 0.01$.

B) Representative western blot images and densitometry quantification showing the expression of Olig1 protein in untransduced and LV.hGALC-transduced ND and GLD NPCs (d0) and differentiated neural progeny (d7, d14 and d24). β -actin was used as loading control. Data are the mean+SEM; n= 2 independent experiments, 1-3 clones/group. Data were analyzed by two-way ANOVA followed by Dunnett's multiple comparison post-test (control group: ND at the corresponding time point). $^*p<0.05$.

### Figure S5

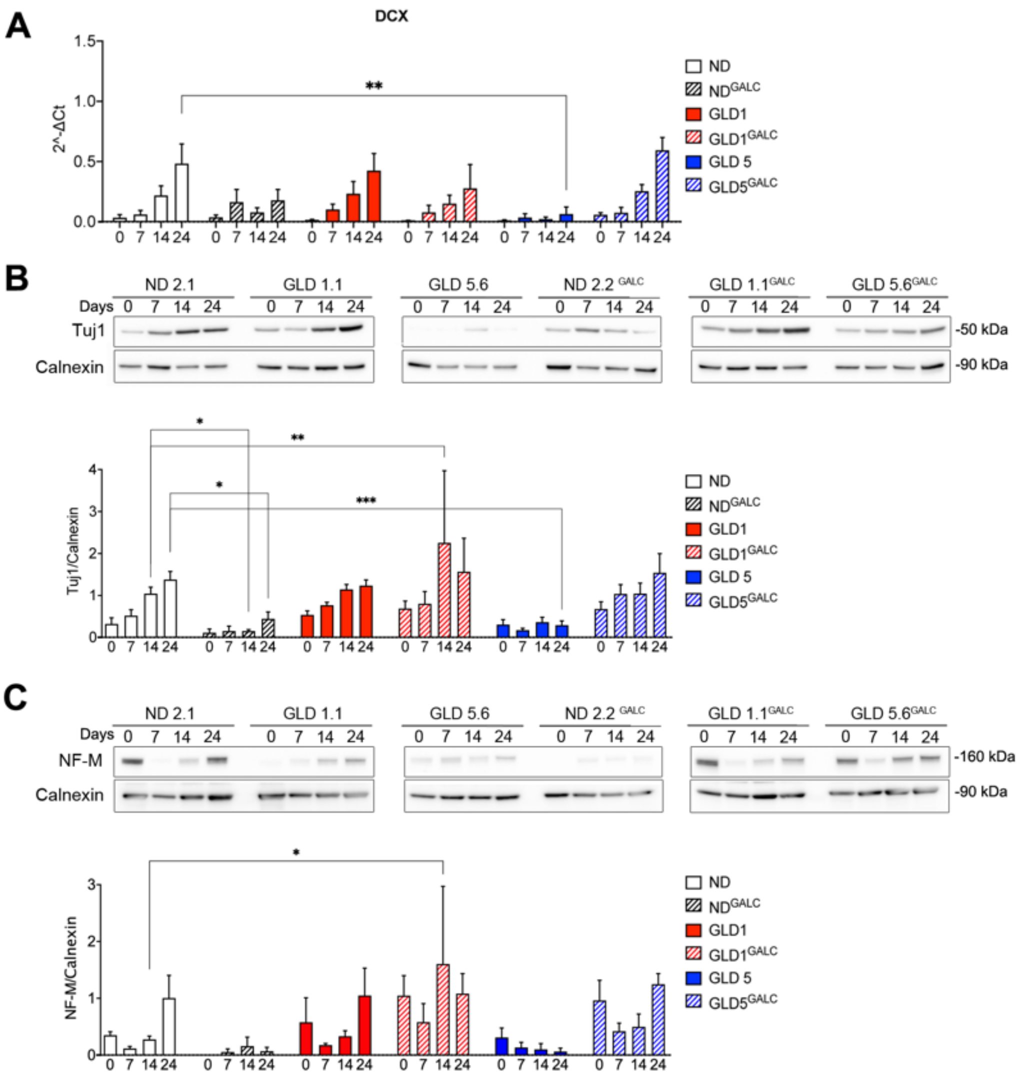

**Figure S5. Expression of neuronal markers during NPC differentiation**

A) Doublecortin (DCX) mRNA expression (assessed by qRT-PCR) in untransduced and LV.hGALC-transduced ND and GLD NPCs (d0) and differentiated neural progeny (d7, d14 and d24). Values are normalized on GAPDH and expressed as 2^-ΔCT (mean+SEM). Clones used: ND1.2, ND2.1, ND2.2, GLD1.1, GLD1.3, GLD5.2, GLD5.6, ND2.2$^{GALC}$, GLD1.1$^{GALC}$, GLD5.2$^{GALC}$, GLD5.6$^{GALC}$. n= 2-4 independent experiments, 1-3 clones/group. Data were analyzed by two-way ANOVA followed by Dunnett's multiple comparison post-test (control group: ND at the corresponding time point). *$p < 0.05$,**$p < 0.01$, ***$p < 0.001$.

B-C) Representative western blot images and densitometry quantification showing the expression of β-tubulin III (Tuj1; B) and Neurofilament Medium (NF-M; C) protein in untransduced and LV.hGALC-transduced ND and GLD NPCs (d0) and differentiated neural progeny (d7, d14 and d24). Calnexin was used as loading control. Data are the mean+SEM; n= 2 independent experiments, 1-3 clones/group. Clones used: ND1.2, ND2.1, ND2.2, GLD1.1, GLD1.3, GLD5.2, GLD5.6, ND2.2$^{GALC}$, GLD1.1$^{GALC}$, GLD5.2$^{GALC}$, GLD5.6$^{GALC}$. Data were analysed by two-way ANOVA followed by Dunnett's multiple comparison post-test (control group: ND at the corresponding time point). *$p<0.05$.

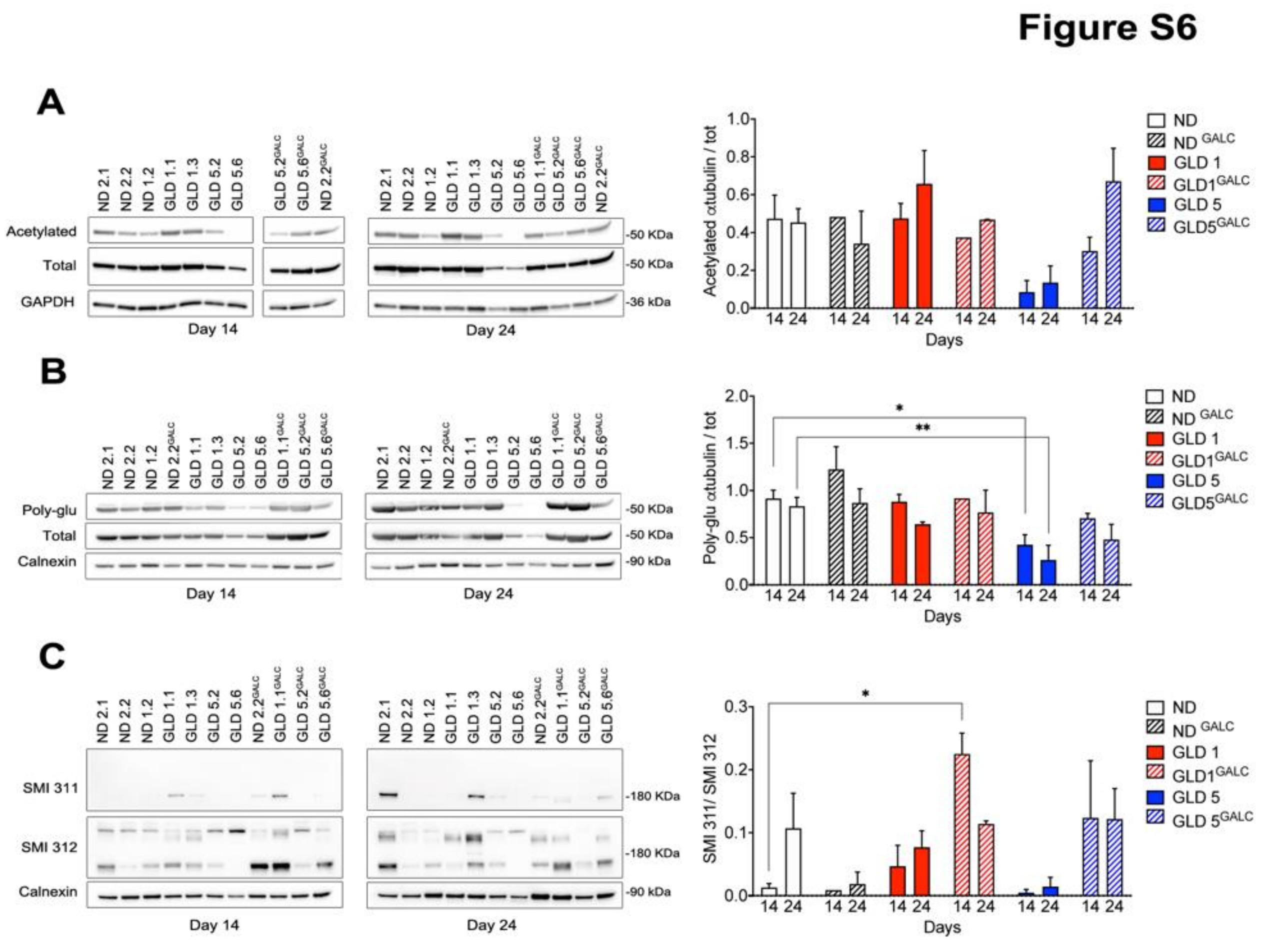

**Figure S6 Expression of neuronal cytoskeleton proteins during NPC differentiation**

(A-C) Representative western blot images and densitometry quantification showing the expression of acetylated tubulin (A), poly-glutamylated tubulin (B) and phosphorylated neurofilaments (SMI311; C) in untransduced and LV.GALC-transduced ND and GLD NPC differentiated neural progeny (d14 and d24). GAPDH or Calnexin were used as loading control. Data were normalized on total tubulin (A, B) and neurofilaments (SMI312; C). Data are the mean+SEM***.*** n= 2 independent experiments, 1-3 clones/group. Data were analysed by two-way ANOVA followed by Dunnett's multiple comparison post-test (control group: ND at the corresponding time point). *p<0.05, **p <0.01.

**Figure S7**

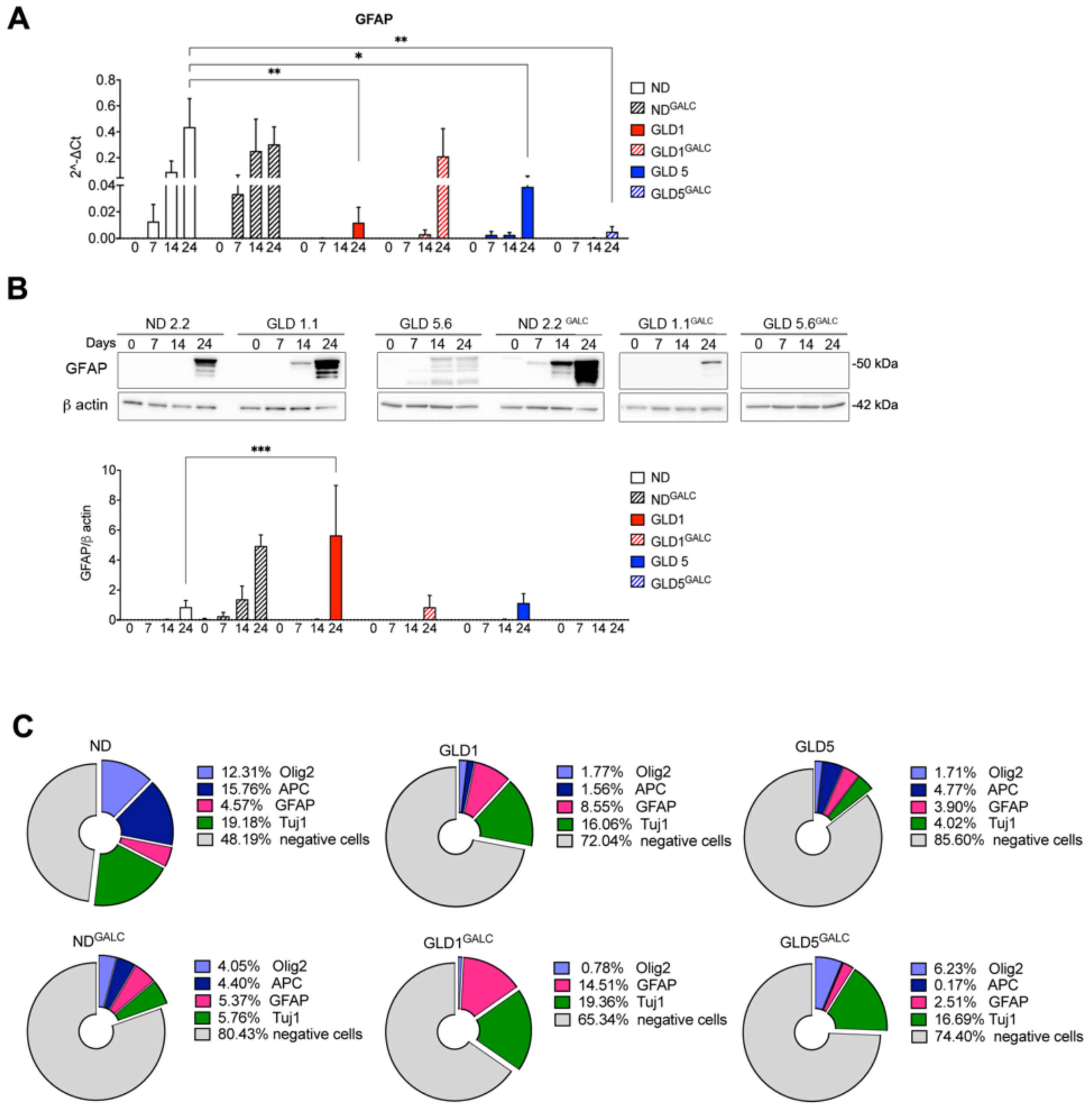

**Figure S7. Expression of astrocytic markers and composition of differentiated cultures**

A) Glial fibrillary acidic protein (GFAP) mRNA expression (assessed by qRT-PCR) in untransduced and LV.hGALC-transduced ND and GLD NPCs (d0) and differentiated neural progeny (d7, d14 and d24). Values are normalized on GAPDH and expressed as 2^-ΔCT (mean+SEM). n= 3-5 independent experiments, 1-3 clones/group. Clones used: ND1.2, ND2.1, ND2.2, GLD1.1, GLD1.3, GLD5.2, GLD5.6, ND2.2$^{GALC}$, GLD1.1$^{GALC}$, GLD5$^{GALC}$, GLD5.2$^{GALC}$, GLD5.6$^{GALC}$. Data were analyzed by two-way ANOVA followed by Dunnett's multiple comparison post-test selecting (control group: ND at the corresponding time point). *$p < 0.05$, **$p < 0.01$, ***$p < 0.001$. Where not indicated, $p>0.05$

B) Representative western blot images and densitometry quantification showing the expression of GFAP protein in untransduced and LV.hGALC-transduced ND and GLD NPCs (d0) and differentiated neural progeny (d7, d14 and d24). β -actin was used as loading control. Data are represented as the mean+SEM. n= 3-5 independent experiments, 1-3 clones/group. Clones used: ND1.2, ND2.1, ND2.2, GLD1.1, GLD1.3, GLD5.2, GLD5.6, ND2.2$^{GALC}$, GLD1.1$^{GALC}$, GLD5$^{GALC}$, GLD5.2$^{GALC}$, GLD5.6$^{GALC}$**.** Data were analyzed by two-way ANOVA followed by Dunnett's multiple comparison post-test (control group: ND at the corresponding time point). ***p < 0.001.

C) Percentages of oligodendrocytes (APC, Olig2+), neurons (Tuj1), and astrocytes (GFAP) in differentiated neuronal/glial cultures (day 24) derived from untransduced (top panel) and LV.hGALC-transduced (bottom panel) ND, GLD1, and GLD5 clones. Data represent the percentage of immunoreactive cells on the total nuclei (mean); n= 2-5 independent experiments; 1-3 clones/group, 5-10 fields/coverslips were analyzed. Clones used: ND1.2, ND2.1, ND2.2, GLD1.1, GLD1.3, GLD5.2, GLD5.6, ND2.2$^{GALC}$, GLD1.1$^{GALC}$, GLD5.2$^{GALC}$, GLD5.6$^{GALC}$.

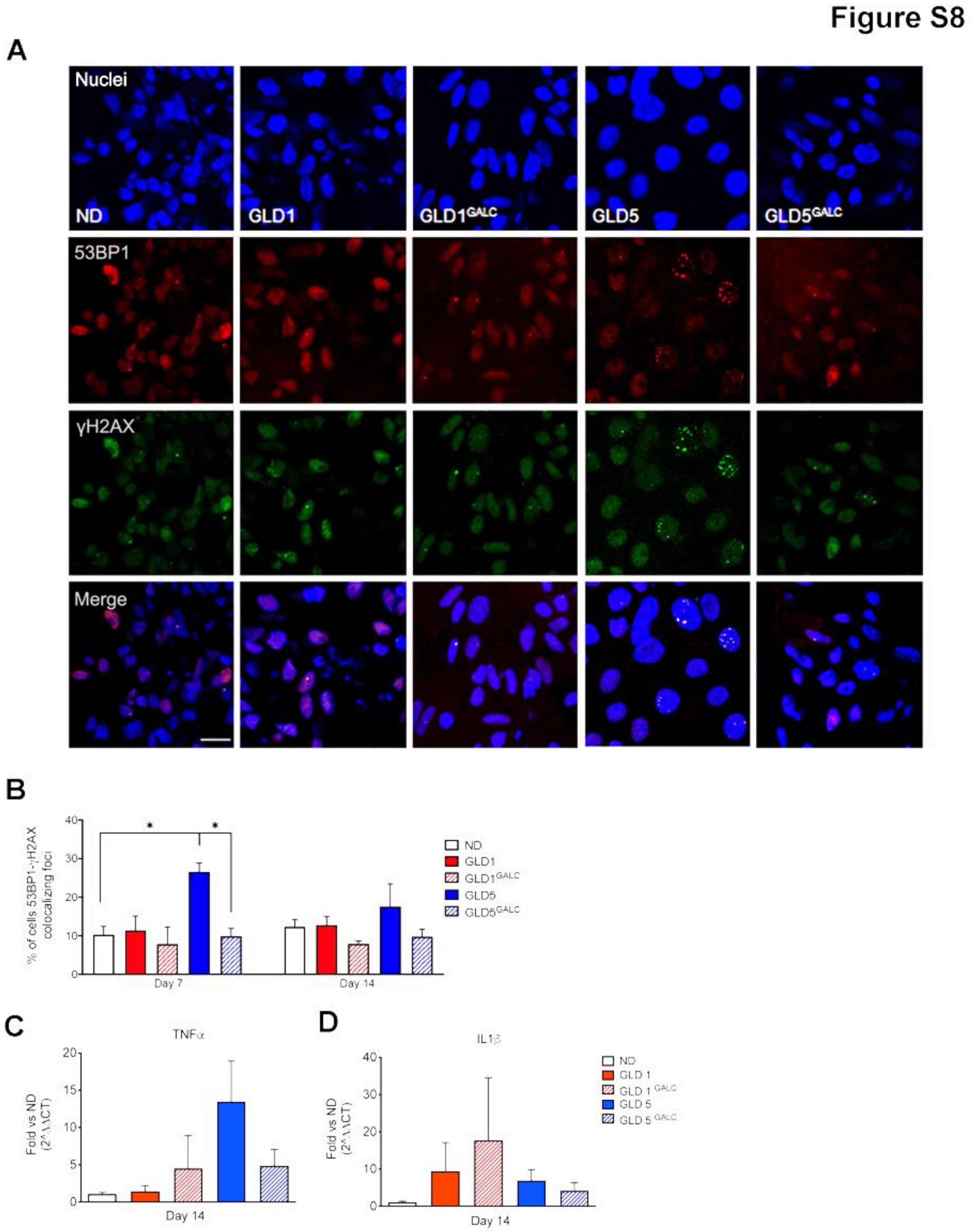

**Figure S8. Expression of senescence markers in ND and GLD NPC-derived cultures**

A-B) Representative pictures (A) and quantification (B) of cells expressing the DNA damage associated markers 53BP1 (red) and γH2Ax (green) in ND, GLD, and $GLD^{GALC}$ NPC (d0) and differentiated cultures (d7 and d14; d14 is shown in the pictures). Nuclei are counterstained with DAPI. Data represent the percentage nuclei with co-localizing 53BP1 and γH2AX foci, and are expressed as the mean + SEM. n= 3 independent experiments, 2-3 replicates/clone for each time point. 100 to 600 nuclei per condition were analyzed. Scale Bar = 15 μm (for all panels, shown in ND). Clones used:

ND1.2, ND2.2, GLD1.1, GLD1.3, GLD5.2, GLD5.6, GLD1.1$^{GALC}$, GLD5.2$^{GALC}$, GLD5.6$^{GALC}$. Data were analyzed by Kruskal Wallis test followed by Dunns' multiple comparison post-test. * $p<0.05$.

C-D) Expression of TNF$\alpha$ (C) and IL1$\beta$ (D) mRNA in ND, GLD, and GLD$^{GALC}$ NPC (d0) and differentiated cultures (d14). n= 2-3 independent experiments; 1-2 clones/group. Clones used: ND1.2, ND2.2, GLD1.1, GLD1.3, GLD5.2, GLD5.6, GLD1.1$^{GALC}$, GLD5.2$^{GALC}$, GLD5.6$^{GALC}$. Data are normalized on the housekeeping gene β-glucuronidase (*GUSB*) and expressed as $2^{-\Delta CT}$. Data are plotted as mean+SEM.

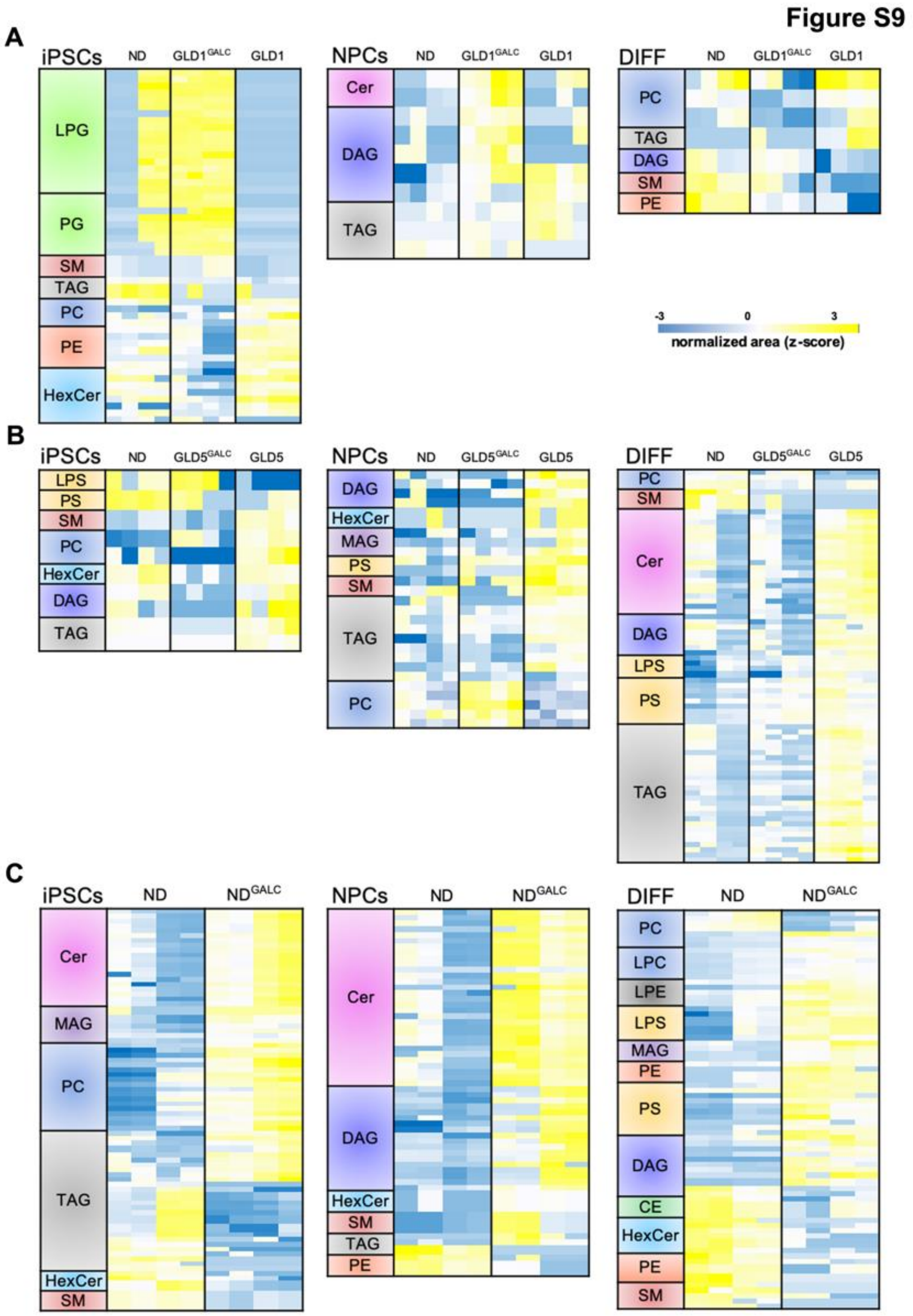

**Figure S9. Lipidomics analysis of untreated and LV.hGALC-transduced ND and GLD cells during iPSC to neural differentiation**

Hierarchical clustering of lipid species differently enriched in iPSCs, NPCs and differentiated cultures (d14) from: A) ND, GLD1, GLD1$^{GALC}$ cells; B) ND, GLD5, GLD5$^{GALC}$ cells; C) ND and ND$^{GALC}$ cells. ANOVA statistical test was applied (FDR<0.05). See Methods for abbreviation of lipid classes.
